# Generation of antagonistic biparatopic anti-CD30 antibody from an agonistic antibody by precise epitope determination and utilization of structural characteristics of CD30 molecule

**DOI:** 10.1101/2024.08.02.606455

**Authors:** Hiroki Akiba, Tomoko Ise, Reiko Satoh, Yasuhiro Abe, Kouhei Tsumoto, Hiroaki Ohno, Haruhiko Kamada, Satoshi Nagata

**Affiliations:** Graduate School of Pharmaceutical Sciences, Kyoto University, Sakyo-ku, Kyoto, 606-8501, Japan; Laboratory of Advanced Biopharmaceuticals, Center for Drug Design Research, National Institutes of Biomedical Innovation, Health and Nutrition, Ibaraki, Osaka, 567-0085, Japan; Laboratory of Antibody Design, Center for Drug Design Research, National Institutes of Biomedical Innovation, Health and Nutrition, Ibaraki, Osaka, 567-0085, Japan; Division of Drugs, National Institute of Health Sciences, 3-25-26, Tonomachi, Kawasaki-ku, Kawasaki, Kanagawa, 210-9501, Japan; School of Engineering, The University of Tokyo, Bunkyo-ku, Tokyo, 113-8656, Japan; Institute of Medical Sciences, The University of Tokyo, Minato-ku, Tokyo, 108-8639, Japan

## Abstract

CD30 is a type I membrane protein that has been successfully targeted for lymphoma therapy using Brentuximab vedotin, an antibody-drug conjugate. Recently, the potential of blocking CD30-dependent NF-κB intracellular signaling has gained attention for treating inflammatory disorders. Development of antibody-based CD30 antagonists would broaden therapeutic strategies. A challenge in developing antagonistic antibodies is that the bivalent form of natural antibody format inevitably cross-links trace amounts of CD30 molecules, leading to signal transduction. In this study, we developed a series of biparatopic antibodies with each pair of antibody variable domains (Fvs) binding to distinct epitopes on CD30, and evaluated their biological activities and binding modes. Initially, we precisely identified epitope sites of the nine antibodies precisely by assessing binding to multiple orthologous CD30 proteins and mutants. We then produced 36 biparatopic antibodies covering all possible combinations of the nine Fvs, and analyzed their biological activities. Among these, we identified both potent agonists and antagonists. Notably, a significant proportion of the biparatopic antibodies displayed reduced agonistic activities, including 1:1-binding antagonists derived from a Fv of a strong agonist previously developed for lymphoma therapy, AC10. The mechanism of signaling activity induction is discussed using epitope information, which leads to the strategies of the development of biparatopic antibodies.

## Introduction

Human CD30 (UniProt ID: P28908) has been pursued as a target of antibody therapy for lymphoma. CD30 was discovered as an overexpressed cell surface molecule in Hodkin’s lymphoma and anaplastic large cell lymphoma. It was later determined to be encoded by member 8 of tumor necrosis factor receptor super family gene (TNFRSF8), with its expression is primarily restricted in activated lymphocytes under normal conditions (1-4), hence its predominant expression in the cancer cells. Consequently, many anti-CD30 monoclonal antibodies (mAbs) had been evaluated over time as antibody therapeutics. The moderate potency of most of the mAbs hindered progress until the approval of Brentuximab vedotin, an antibody-drug conjugate (ADC) successfully introduced in the market (5). Besides the potent anti-mitotic activity of vedotin drug moiety, Brentuximab vedotin utilizes an antibody known as AC10, originally developed as an agonist as naked antibody. AC10 initiates CD30-mediated signaling including activation of NF-κB pathway, which may contribute on controlling tumor cell growth. Despite the initial expectation that agonists targeting CD30 would induce apoptosis signaling in expressing cells, clinical outcomes, including those with AC10, failed to demonstrate significant advantages (6,7). Antagonists targeting CD30 have not yet been developed.

In natural conditions, activation of CD30 occurs through binding with its trimeric ligand, CD153 (CD30L). Members of the TNFRSF, such as CD30, contain variable numbers of cysteine-rich domains (CRDs) in their extracellular regions, which form their ligand binding sites. CRDs, commonly observed in >20 TNFRSF members, share a common fundamental high-ordered structure (8,9). The ligand molecules of the tumor necrosis factor superfamily, including CD153, are trimeric and induce clustering of corresponding TNFRSF on the cell membrane through their interaction (Fig. 1A) (1,10). The reassembled TNFRSF into cluster configurations involves conformational changes in the intracellular regions at a macromolecular level, and downstream signaling molecules such as TNFR-associated factors are recruited to activate NF-κB pathway.

**Fig.1.**
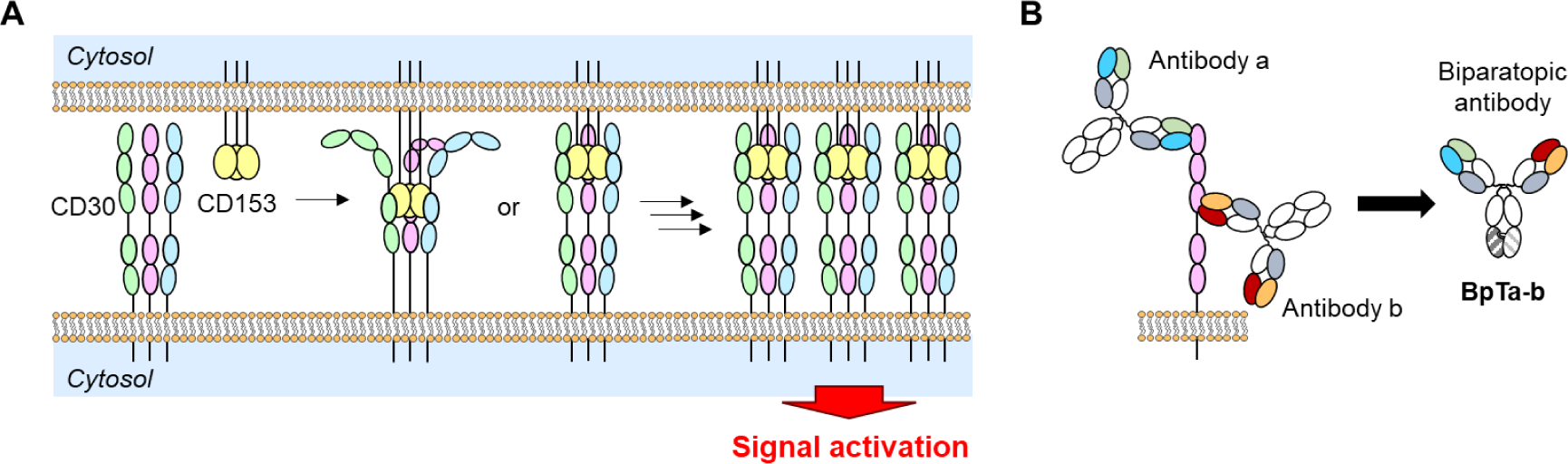
A) Activation mechanism of CD30. CD30 has two regions binding the specific trimeric ligand (CD153). With the association of CD153, CD30 is expected to form clusters on the cell membrane, and downstream signal cascade is activated. B) Bivalent biparatopic antibodies (BpAbs). In this study, BpAbs are named as BpTa-b after two originating antibodies (antibody a and antibody b).

It has been well established that TNFRSF signaling plays crucial and diverse immunomodulatory roles in various pathological conditions, including a range of immune pathologies, cancers, and infectious diseases (11-13). Given the frequent association of the CD30-CD153 axis with immunological disorders (14-18), antagonists are anticipated as drug candidates to mitigate inflammatory responses. Although agonistic or antagonistic activities of various anti-CD30 monoclonal antibodies are anticipated to differ in an epitope-dependent fashion, the significance of the epitope location remains elusive because the previous structure-function analyses were complicated by the unusual structure of human CD30 containing large duplicate regions with two predicted ligand binding sites. This has prevented rational design and selection both of agonists and antagonists. Additional challenge in developing antagonistic antibodies is the bivalency of natural antagonistic antibody inevitably cross-links trace amounts of target CD30 molecules, leading to signal transduction.

Herein, we focus on biparatopic antibodies (BpAbs), which are engineered bispecific antibodies using two distinct variable fragments (Fv) to two different epitopes of a single antigen molecule (Fig. 1B) (19,20). Recently, we successfully produced an antagonist against tumor necrosis factor receptor 2 (TNFR2), another member of the TNFRSF, using BpAb technology (21). Similar to CD30, TNFR2 is activated by clustering upon binding to trimeric TNFα ligand. While bivalent binding by conventional bivalent antagonistic antibodies induced a moderate level of signaling, the produced anti-TNFR2 BpAb antagonist bound TNFR2 in a 1:1 manner, resulting in no signal induction and maximizing antagonistic activity (21). Building on this finding, we hypothesized that antagonists against TNFRSF, including CD30, could be developed using a similar approach.

In this study, we selected a panel of nine previously-made anti-CD30 mAbs whose epitopes are distributed across the six CRDs on the extracellular domain of the human CD30 molecule (5,22-25). Since each pair of the nine antibodies recognize a unique pair of topographical epitopes on CD30 molecules, we developed the biparatopic antibody series using all possible pairs of the nine antibody variable regions. We comprehensively evaluated the relationship of binding mode and CD30 downstream signaling potency. We identified both potent agonists and antagonists. A significant proportion of the BpAbs exhibited reduced agonistic activities. Antagonistic BpAbs were identified, including those employing Fv from an agonist antibody in one arm, such as AC10. We discuss the insight obtained through the characterization of the binding modes in the contexts of the possible mechanism responsible for the signaling activities. In particular, we explore the molecular mechanisms underlying the conversion of agonists to antagonists. The overall data suggest that the targeted epitope can be selected for BpAbs with desired functionality with different mode of actions for improved therapeutic activity.

## Results

### Nine topographical epitope groups recognized by the panel of selected nine anti-CD30 mAbs

We selected nine anti-human CD30 antibodies for this study from a pool of previously known 42 antibodies. The nine mAbs were generated in mice and are capable of reacting with distinct topographical epitopes which were determined by mutual competition assays. The nine epitopes span across the native conformational structure of the extracellular domain of human CD30 (22-25). Each topographical epitope encompasses multiple residue spans within the modular structure of CD30 (25,26). The names of the mAbs are T104, T6, T107, T427, T426, AC10, T105, T25, and T405. Among them, AC10, is the original antibody comprising the ADC brentuximab vedotin (5). Epitope information of other antibodies unused in this study is summarized based on literatures (Supplemental Text).

The antibodies were made as human IgG1κ chimera antibodies in conventional human IgG1 chimera format (cAb). Competitive binding of the cAbs was first analyzed in an enzyme-linked immunosorbent assay (ELISA) using CD30 fused to rabbit Fc (CD30-rFc) and by surface plasmon resonance (SPR) using maltose-binding protein-fused CD30 (CD30-MBP; Fig. S1). In ELISA-based assay (Table 1), binding of CD30-rFc to the indicator antibody (y-axis) in the presence of competitor (x-axis). In SPR analysis (Table 2 and Fig. S2), CD30-MBP was flowed onto immobilized first antibody as cAb and then second antibody as F(ab′)_2_ fragment was flowed. There was slight difference among duplicate-binding antibodies (T6, T107 and T427) (25), probably due to the use of dimeric antigen in the case of ELISA. For both cases, T426–T105, AC10–T427, and T427–T107 pairs showed complete competition. These results were consistent with partly reported previous results (24). Relative epitope positions were not influenced by rebuilding of antibodies into cAbs.

**Table 1.** Mutual competition of cAbs by enzyme-linked immunosorbent assays.

|  |  | Indicator cAb |  |  |  |  |  |  |  |  |
| --- | --- | --- | --- | --- | --- | --- | --- | --- | --- | --- |
|  |  | T104 | T426 | AC10 | T105 | T25 | T405 | T6 | T427 | T107 |
| Competitor<br>cAb | T104 | 11 | 91 | 93 | 94 | 92 | 93 | 95 | 94 | 93 |
|  | T426 | 138 | 13 | 89 | 3 | 145 | 156 | 133 | 119 | 155 |
|  | AC10 | 107 | 97 | 10 | 101 | 101 | 101 | 22 | 22 | 38 |
|  | T105 | 97 | 46 | 87 | 0 | 95 | 97 | 86 | 91 | 95 |
|  | T25 | 104 | 102 | 101 | 104 | 10 | 95 | 25 | 25 | 76 |
|  | T405 | 106 | 103 | 101 | 108 | 93 | 11 | 106 | 104 | 101 |
|  | T6 | 103 | 102 | 83 | 99 | 88 | 98 | 19 | 20 | 36 |
|  | T427 | 101 | 94 | 85 | 99 | 94 | 97 | 18 | 19 | 36 |
|  | T107 | 104 | 102 | 83 | 99 | 80 | 100 | 28 | 32 | 28 |

**Table 2.** Mutual competition of cAbs by surface plasmon resonance^a^.

|  |  | Second antibody, F(ab')<sub>2</sub> |  |  |  |  |  |  |  |  |
| --- | --- | --- | --- | --- | --- | --- | --- | --- | --- | --- |
|  |  | T104F | T426F | AC10F | T105F | T25F | T405F | T6F | T427F | T107F |
| First antibody, cAb | T104 |  | x | x | x | x | x | x | x | x |
|  | T426 | x |  | x |  | x | x | x | x | x |
|  | AC10 | x | x |  | x | x | x | x |  | x |
|  | T105 | x |  | x |  | x | x | x | x | x |
|  | T25 | x | x | x | x |  | x | x | x | x |
|  | T405 | x | x | x | x | x |  | x | x | x |
|  | T6 | x | x | x | x | x | x |  | x | x |
|  | T427 | x | x |  | x |  | x | x |  | - |
|  | T107 | x | x |  | x | x | x | x |  |  |

<sup>a</sup> x and - represent no competition and partial competition, respectively.

### Ortholog screening for epitope determination

Human CD30 is an unusual member of the TNFRSF with a duplicated segment within its extracellular domain that may function as an additional ligand binding site. The extracellular region of human CD30 spans approximately 360 amino acids and contains six cysteine-rich domains (CRDs), commonly bound by specific ligands of the TNFRSF (Fig. 2A). Regions between CRD3 and CRD5, including the entire CRD4, are predicted to be disordered according to AlphaFold2 (Fig. 2B) (27,28). High homology regions exist within CRD1-3 and CRD4-6, likely due to gene duplication (Fig. 2C,D). Our experiments confirmed that these regions also exhibit functional similarity, with a common binding ability to CD153, the specific ligand for CD30 (Fig. 2E).

**Fig. 2.**
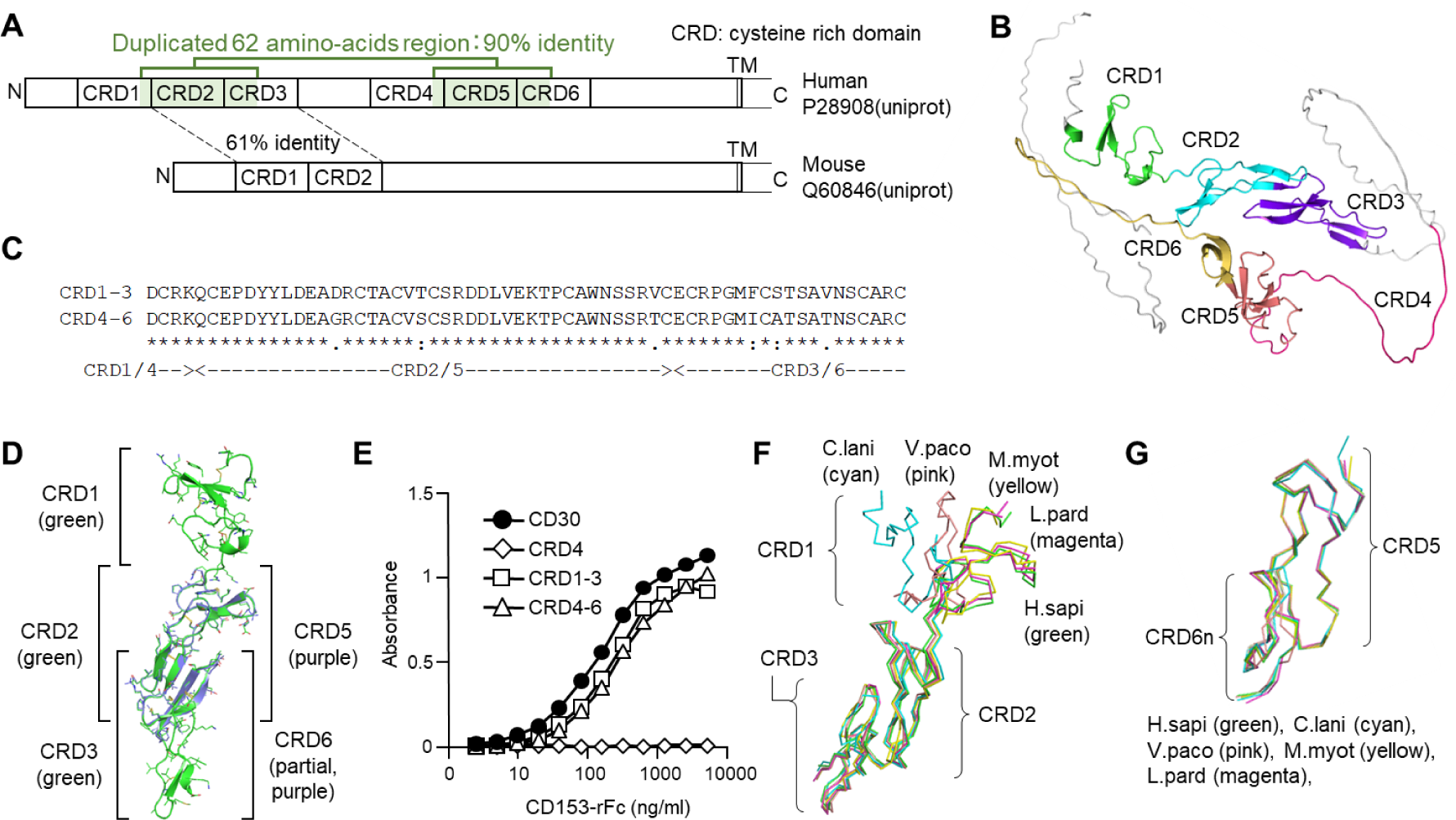
Structural characteristics of human CD30. A) Schematic illustration of human CD30 with duplicate and ligand-binding regions. B) Presence of disordered regions. In addition to the region between CRD3 and CRD4, CRD4 (magenta) and C-terminal half of CRD6 are predicted to be disordered in AlphaFold2. C) Similarity of CRD2-3 and CRD5-6. D) Superimposed AlphaFold-predicted structures of CRD1-3 and CRD5-6n from human CD30. E) Binding of CD153 both to CRD1-3 and CRD4-6 recombinant proteins. F) Superimposed AlphaFold-predicted structures of CRD5-6n from human and five orthologs. G) Superimposed AlphaFold-predicted structures of CRD1-3 from human and five orthologs.

We initially aimed at precisely identifying amino acids residues associated with the epitope structures of the anti-CD30 antibodies. As reported earlier for another TNFRSF member, TNFR2, precise determination of epitope sites (down to approximately 2–10 amino acids) was necessary for mechanistic analysis and the design of BpAbs with biological activities (21). In the previous study, antibody epitopes were determined using mutants substituting human TNFR2 with mouse TNFR2 sequences (21). Despite the similarity, the latter’s structure was immune-tolerated in the mouse species used for mAb production. This resulted in the cancellation of specific mAb binding if the substituted part hits the epitope location. In the case of CD30 in this study, similar epitope determination strategy based on human/mouse substitution posed challenges due to the duplicated regions because mouse CD30 (Uniprot ID: Q60846) possesses three CRDs only and substitutions are not logically designed due to the absence of this duplicate region (29).

To address this issue, ortholog genes of CD30 were screened for similarity to human CD30. A BLAST search of the protein sequence of the extracellular region of human CD30 was conducted (30), and ortholog sequences showing 45–65% identity were selected. To ensure diversity among the orthologs, five sequences, spaced apart in a phylogenetic tree, were chosen (Fig. S3). These sequences correspond to predicted *tnfrsf8* gene products from *Tupaia chinensis* (*Tc*; Refseq ELW68632.1), *Chinchilla lanigera* (*Cl*; Refseq XP_013359778.1), *Lynx pardinus* (*Lp*; Refseq VFV35557.1), *Vicugna pacos* (*Vp*; Refseq XP_006197082.2), and *Myotis myotis* (*Mm*; Refseq KAF6380682.1) (amino acid sequences in Table S1). Upon transient expression of the five ortholog *tnfrsf8* genes in HEK293T cells, all five gene products were found to be bound by at least one of the nine antibodies (Fig. 3A), indicating expression of functional proteins on the cell membrane. Additionally, total or partial reduction in antibody binding level was observed for each *tnfrsf8* gene product.

**Fig. 3.**
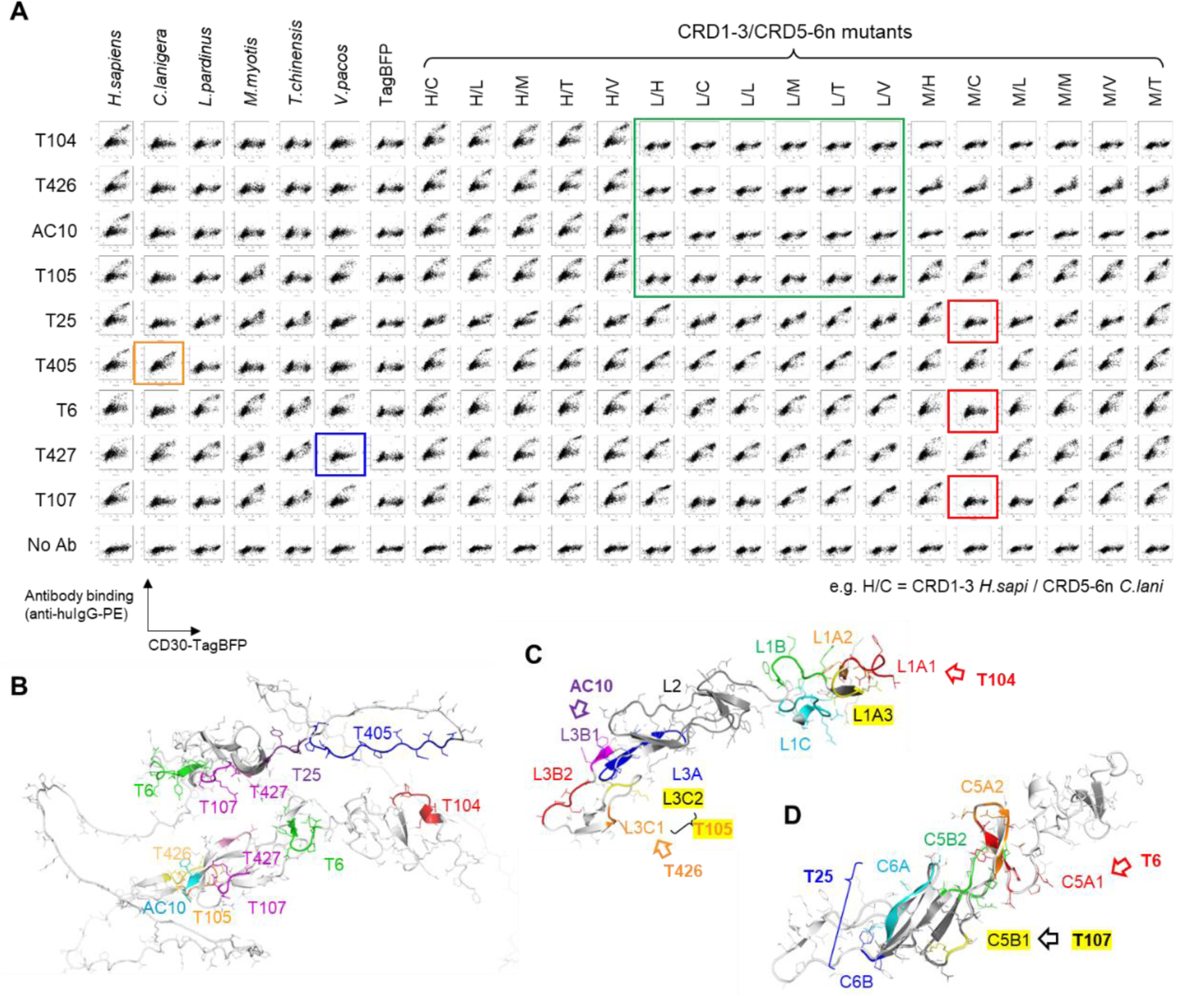
Epitope determination using CD30 mutants. A) Affinity reduction of the antibodies with multi-domain substituted CD30 analyzed in flow cytometer. Green, red, blue, and orange boxes are key mutants or orthologs for epitope determination in the initial step. B) Determined epitope sites of all antibodies. C) CRD1-3 mutants used to determine epitopes of T104, AC10, T105, and T426. D) CRD5-6n mutants used to determine epitopes of T6, T25, and T107.

Next, structures of the CRDs of the ortholog *tnfrsf8* gene products were predicted using AlphaFold2 (27). This allowed for a focus on structurally similar protein domains suitable for substitution. CRD1, CRD2, CRD3, and CRD5 of human CD30 exhibit the typical CRD textually folding structure observed in TNFRSF proteins. In addition, the N-terminal half of CRD6 is folded. The structural similarity directly relates to protein function and is more robust than random amino acid sequence changes, which may disrupt folding structures necessary for accurately determining conformational epitopes. Similar predicted structures were found for the five *tnfrsf8* gene products from the orthologs, with common features including disordered CRD4 and the C-terminal half of CRD6. CRD1-3 of *Tc* are likely disordered, and CRD1 of *Cl* and *Vp* may also exhibit a different fold from CRD1 of humans (Fig. 2F). Structures of CRD5 and the N-terminal half of CRD6 (CRD5-6n) were predicted to be similar among the orthologs (Fig. 2G). Utilizing this information, CRD1-3 was substituted with those from *Lp* and *Mm*, while CRD5-6n was substituted with those from all five orthologs. In total, including human CD30, 3 × 6 = 18 clones were produced (Table S1), and their binding to these substituted mutants was analyzed by flow cytometry. The results revealed several patterns of affinity reduction (Fig. 3A), and further optimization of the mutants was conducted based on the antibody-dependent reduction pattern.

### Epitope determination

Epitope determination was conducted using partially substituted CD30 expressed on the cell membrane. All mutants used in this study are listed in Table S1. The determined epitope regions are listed and mapped on the AF2-predicted structure of human CD30 (Table 3, Fig. 3B). Among the nine antibodies, three (T6, T107, and T427) were binders against the duplicate region. Below are the descriptions of epitope determination for each antibody.

*(1) CRD1-3 binders, including T104, AC10, T426, and T105.* These four antibodies showed a common binding pattern independent of CRD5-6n (Fig. 3A). They exclusively recognized CRD1-3. Since all antibodies failed to bind to *Lp*CRD1-3 (green box in Fig. 3A), the critical amino acid residues for constituting the epitope structures were narrowed down by partial substitution to *Lp* sequence. Initially, each CRD was substituted. T104 did not bind to the CRD1 mutant of *Lp* (denoted as L1), while the other three antibodies did not bind to the CRD3 mutant of *Lp* (L3) (Fig. S4). For T104, CRD1 was further split into three based on disulfide connectivity (L1A, L1B, and L1C), and L1A was not bound (Fig. S5). By further narrowing down the mutated region (L1A1, L1A2, and L1A3), a reduction in binding to L1A1 was observed, and this region was assigned as the critical amino acids for the epitope structure (Fig. S5). In a similar strategy, CRD3 was split into three (L3A, L3B, and L3C), and then into two pairs (L3B1 and L3B2, L3C1, and L3C2). Binding was reduced for L3B1 by AC10, and L3C1 by T426 (Fig. S6). Binding of T105 was not observed for L3C, but the split mutants were bound; thus, both regions corresponding to L3C1 and L3C2 contributed partly to the binding of T105 (Fig. 3C, Fig. S6).
*(2) T6, T107 and T25.* These antibodies exhibited reduced binding by *Mm*CRD1-3 and *Cl*CRD5-6n doubly substituted clone (red boxes in Fig. 3A). Based on this, a partial *Cl* mutant was introduced to CRD5 or CRD6 of *Mm*CRD1-3, and a similar method was conducted as in (1) to narrow down the epitopes. As a result, T6 showed reduced binding to the partial mutant of CRD5 (MC5A1), and T107 similarly showed reduced binding to MC5B1. T25, on the other hand, exhibited only a partial reduction in binding to MC6A. When the adjacent region was also substituted with *Cl* sequence (denoted as MC6B), a reduction in the affinity was observed (Fig. 3D, Fig. S7). Therefore, this region was determined as the epitope.
*(3) T427.* The epitope of T427 was previously reported to be inside the duplicate region (25). Among the tested ortholog clones, only *Vp* showed reduced binding (blue box in Fig. 3A). When all CRDs in this region of human CD30 were substituted with the *Vp* sequence (V2356), no binding was observed (Fig. S8). By comparing the peptide sequences to search for regions with multiple amino acid differences between *Vp* and human CRD2-3 and CRD5-6, three pairs of regions were found (Fig. S8). These regions inside CRD2-3 of V2356 were back-mutated to the human sequence (named V2356-EAD, V2356-SRD, V2356-SAV). The SRD mutation fully restored binding, the SAV mutation partially, while the EAD mutation did not restore binding (Fig. S8). Thus, S88-D90/S263-D265 was found as the epitope core and S117-N120/S292-N295 in the periphery.
*(4) T405.* The epitope of T405 was reported to be inside CRD6 (25). Among the ortholog sequences, only *Cl* showed binding (orange box in Fig. 3A), thus, the region that only *Cl* shows high homology in the C-terminal disordered region of CRD6 was the candidate. This coincided with D321-C335 (Fig. S9). When these regions were substituted with *Mm* sequence (M6e), or split in half (M6e1 and M6e2), all showed reduced binding (Fig. S9). Thus, the epitope of T405 was determined to be this whole region.

**Table 3.**
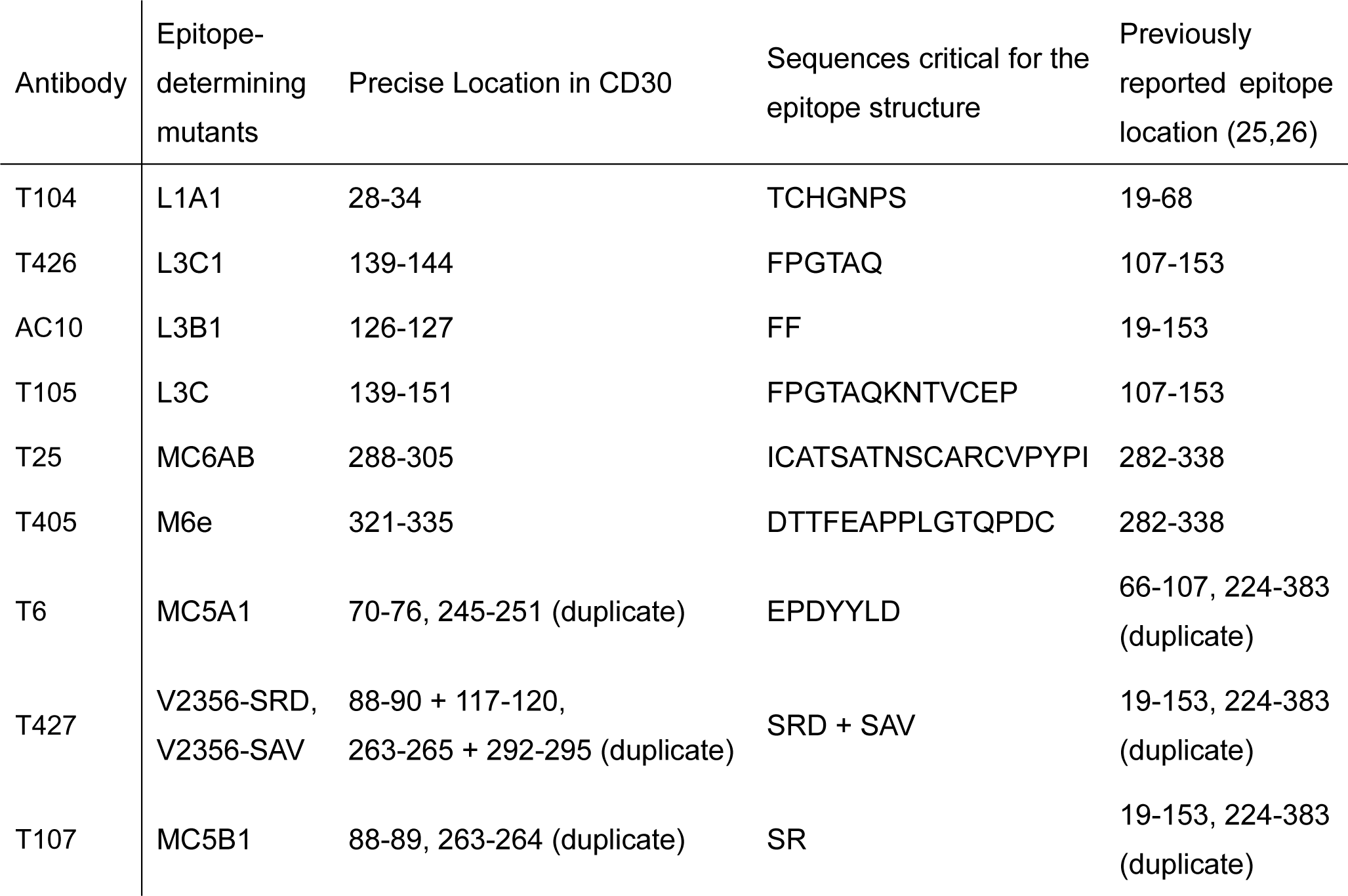
Epitopes of the antibodies determined by mutant analysis.

### Production and binding activities of the BpAbs

Thirty-six BpAbs were produced from nine Fvs using a previously reported method to cover all biparatopic combinations of epitopes (Table 4) (31). This method utilizes intein-mediated protein trans-splicing (IMPTS) between separately expressed and purified antibody fragments fused either with Int^N^ or Int^C^. The fusion proteins Int^N^- or Int^C^-Fc with antigen-binding fragment (Fab) of each antibody is named with its name in a way such as T104^N^ or AC10^C^. The expression yield of each fragment varied over 6 – 367 mg/L (Table S2). Although few fragments such as T6^N^ showed low expression yield in the Expi293 expression system, probably due to the poor folding ability of each Fv, IMPTS enabled homogeneous BpAb yield independent of the Fv used. The resulting BpAb by IMPTS is named after the identification numbers of the two Fvs, for example, BpT104-10 when T104^N^ and AC10^C^ are used (Table 4). The production method enabled high symmetry except for two amino-acid substitutions in the hinge region of the ‘N’ side Fab.

**Table 4.**
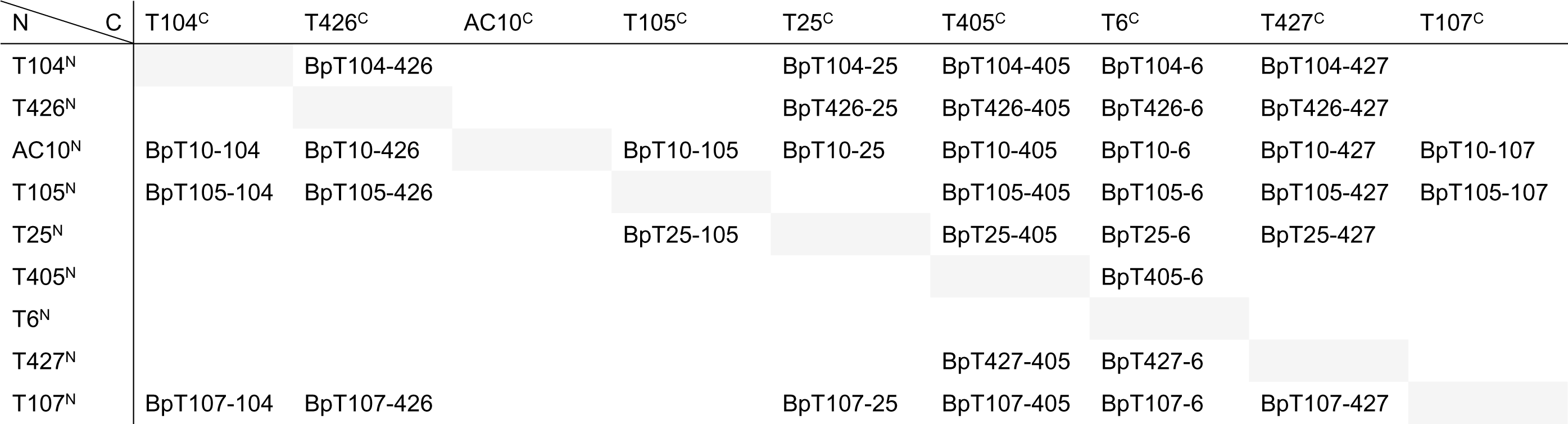
Production of anti-CD30 BpAbs by the intein-fused materials used (such as T104^N^ and T426^C^)

| N \ C | T104<sup>C</sup> | T426<sup>C</sup> | AC10<sup>C</sup> | T105<sup>C</sup> | T25<sup>C</sup> | T405<sup>C</sup> | T6<sup>C</sup> | T427<sup>C</sup> | T107<sup>C</sup> |
| --- | --- | --- | --- | --- | --- | --- | --- | --- | --- |
| T104<sup>N</sup> |  | BpT104-426 |  |  | BpT104-25 | BpT104-405 | BpT104-6 | BpT104-427 |  |
| T426<sup>N</sup> |  |  |  |  | BpT426-25 | BpT426-405 | BpT426-6 | BpT426-427 |  |
| AC10<sup>N</sup> | BpT10-104 | BpT10-426 |  | BpT10-105 | BpT10-25 | BpT10-405 | BpT10-6 | BpT10-427 | BpT10-107 |
| T105<sup>N</sup> | BpT105-104 | BpT105-426 |  |  |  | BpT105-405 | BpT105-6 | BpT105-427 | BpT105-107 |
| T25<sup>N</sup> |  |  |  | BpT25-105 |  | BpT25-405 | BpT25-6 | BpT25-427 |  |
| T405<sup>N</sup> |  |  |  |  |  |  | BpT405-6 |  |  |
| T6<sup>N</sup> |  |  |  |  |  |  |  |  |  |
| T427<sup>N</sup> |  |  |  |  |  | BpT427-405 | BpT427-6 |  |  |
| T107<sup>N</sup> | BpT107-104 | BpT107-426 |  |  | BpT107-25 | BpT107-405 | BpT107-6 | BpT107-427 |  |

Binding of the antibodies to the CD30-stably expressed NF-κB reporter Ramos-Blue cell line and the CD30-expressing lymphoma cell line KARPAS 299 (32) was analyzed. All cAbs and BpAbs showed binding to the CD30-Ranos-Blue cells in a dose-dependent manner (Fig. S10). The reactivities of all cAbs and BpAbs were also conformed with native CD30 molecule expressed on KARPAS 299 cell line (Fig. S11). The affinity of BpAbs as well as the original cAbs was analyzed in surface plasmon resonance (SPR) in rough level. In this experiment, antibodies were first captured on the sensor chip, and CD30-MBP was flowed next. Thus, the effect of avidity by cross-linking pattern was minimized. As the result, all cAbs and BpAbs bound CD30-MBP, but the affinity of the T426 cAb was too weak to determine the parameters correctly (Fig. S12, Table S3). BpAbs including T426-based ones conferred strong affinity to CD30-MBP as typically observed for BpAbs (19). The result indicated that both of Fvs comprising BpAbs play roles in the binding.

### Biological activities of the BpAbs

The biological activities of the cAbs and BpAbs were analyzed using a Ramos-blue based reporter cell line. Expression of CD153, normally observed in Ramos cells, was downregulated in the stable cell line (31). Agonistic activity was analyzed by incubating cells with the antibodies in dilution series, whereas antagonistic activity was analyzed in the presence of CD153-Fc fusion protein. The results are presented in Fig. 4. It was demonstrated that all nine cAbs showed moderate to high levels of agonistic activities. Among them, AC10 showed the strongest maximum agonistic activity reached, and another original cAb, T104, recognizing non-ligand binding site CRD1 showed a similar level of agonistic activity. The maximum activity of other cAbs was lower in the concentration range analyzed. On the other hand, only a small number of cAbs showed antagonistic activity. The most potent antagonist was T427, but a significant level of residual signaling activity was observed. Because T427 showed moderate agonistic activity in the absence of CD153-Fc, the competing activity may have reflected this observation as we observed for TNFR2 in a previous study (21).

**Fig. 4.**
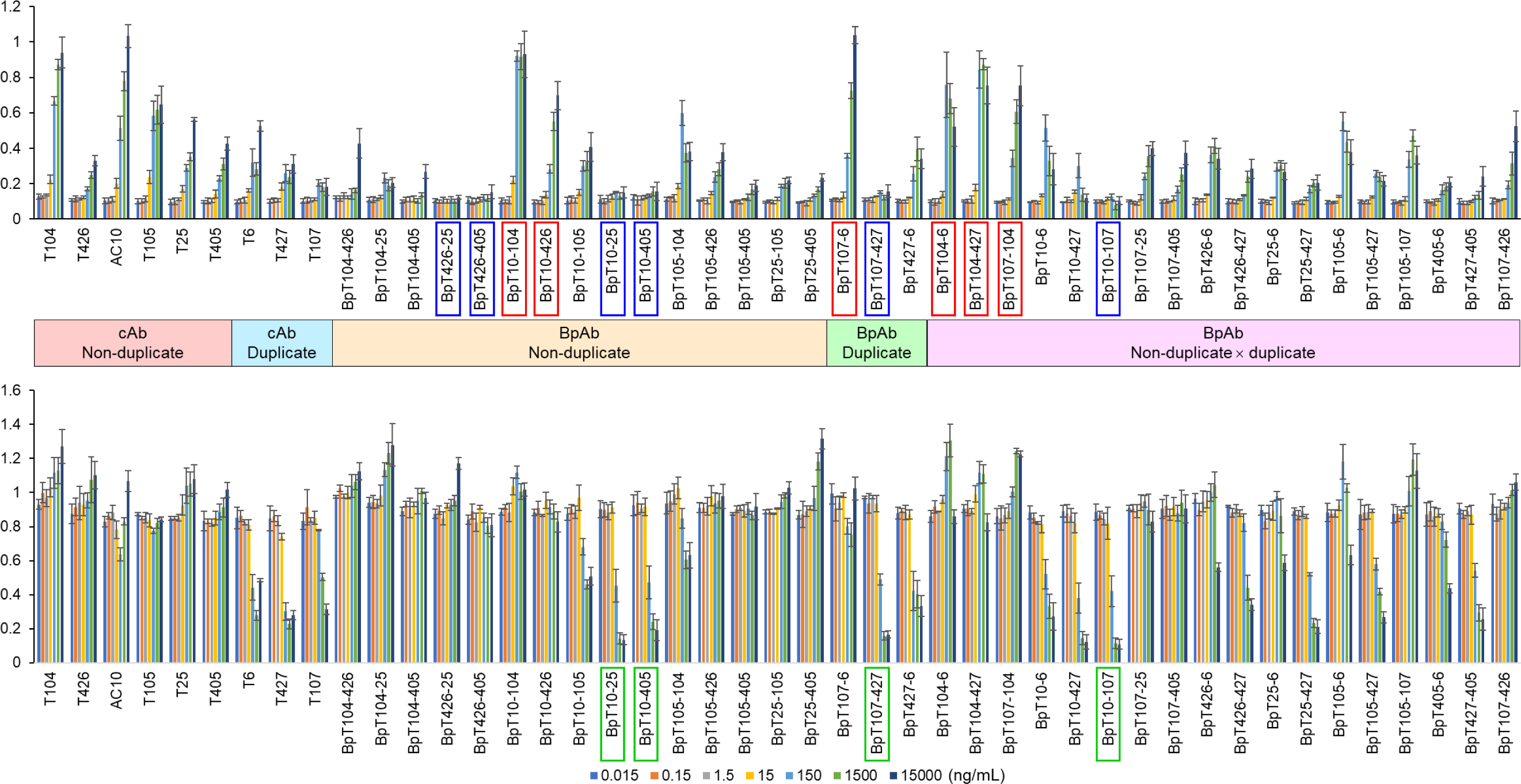
Agonistic and antagonistic activities of the cAbs and BpAbs analyzed using CD30-transfected NF-κB reporter cells. Agonistic (top) and antagonistic (bottom) activities were measured in the absence or presence of CD153-Fc. Antibodies are categorized by their recognizing non-duplicate and duplicate epitopes. Boxed labels indicate strong agonists (red), non-agonists (blue) and effective antagonists (green) among BpAbs.

BpAbs showed different profiles of activities. Six BpAbs (BpT10-104, BpT10-426, BpT107-6, BpT104-6, BpT104-427, and BpT107-104; highlighted in red, Fig. 4, top) showed maximum reached signaling activity similar to AC10. As an agonist, BpT10-104 was superior to AC10 in that a lower concentration was needed to achieve this activity. Interestingly, BpT107-6 was a strong agonist even though the original cAbs (T107 and T6) were not. However, overall, for activating CD30, BpAb offers only a limited level of advance compared to cAbs.

In contrast, improved antagonist candidates were generated among BpAbs. Many BpAbs showed a limited level of agonistic activity. Six BpAbs (BpT426-25, BpT426-405, BpT10-25, BpT10-405, BpT107-427, and BpT10-107; highlighted in blue, Fig. 4, top) showed no agonistic activity. This elimination of agnostic activity of BpAb format led to a number of BpAbs that showed strong antagonistic activities. Among cAbs, T427 and T107 were potential antagonists. Although T6 also showed antagonistic activity, activation of CD30 may compensate for the activity when used in high concentration. BpAbs including BpT10-25, BpT10-405, BpT10-427, BpT107-427, and BpT10-107 (highlighted in green, Fig. 4, bottom) showed stronger antagonistic functions, and apparent signaling activity was not observed. Interestingly, most BpAbs using AC10 Fvs were antagonists. Therefore, the best agonistic cAb was flipped into one of the best antagonist candidates. To elucidate the mechanism of antagonistic activities, binding inhibition of CD153 ligand to the reporter cell line was analyzed in a flow cytometer (Fig. S13). As a result, most antagonists were binding inhibitors of CD153 (Fig. 5). AC10 also was a binding inhibitor of CD153, and thus, functionally hindered potential (ligand-blocking antagonism) arose as a function by building into BpAbs. Reduction of agonistic activity by BpAb generation was the key to the antagonistic function.

**Fig. 5.**
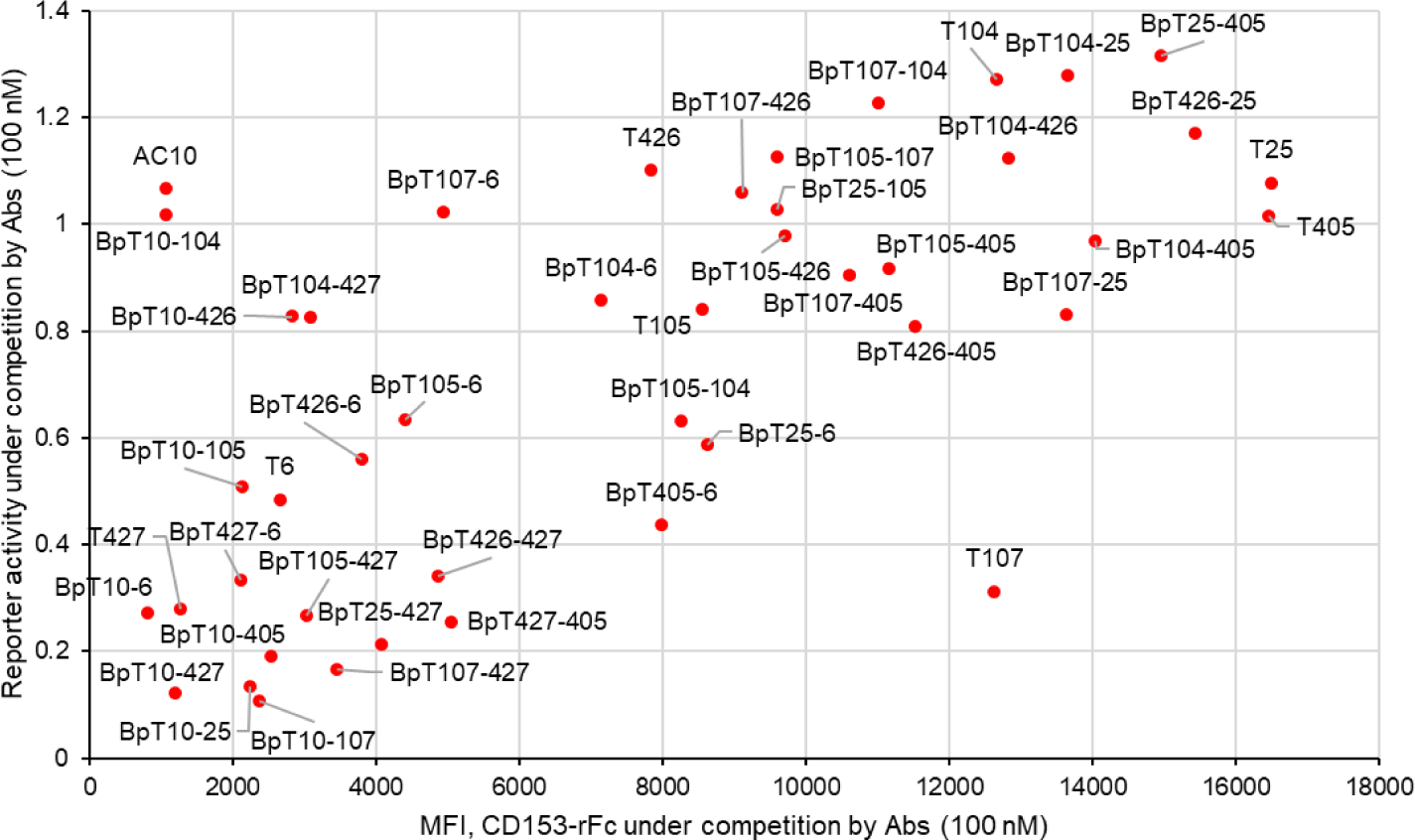
Relationship between the biological antagonistic activity of the cAbs and BpAbs (y-axis) with CD153-binding competition to the CD30-expressing cells (x-axis).

### Size of immunocomplexes

In a previous study, we developed anti-TNFR2 BpAbs that exhibited agonistic activities dependent on the size of immunocomplexes formed in solution with the soluble form of recombinant antigen proteins (21). Using a similar method, we analyzed the size of the immunocomplexes for agonistic and non-agonistic BpAbs using size-exclusion chromatography-multi-angle light scattering (SEC-MALS) (Fig. 6, Fig. S14). To simplify the analysis, two BpAbs, BpT10-104 and BpT10-405, were selected that possess two Fvs binding non-duplicate epitopes. The results showed that the agonist, BpT10-104, formed large immunocomplexes with the majority of 2:2 complexes (Fig. 6A, 12.3 mL peak, approximately 430 kDa), while the non-agonist, BpT10-405, only formed 1:1 immunocomplexes (Fig. 6B, 14.6 mL peak, approximately 220 kDa). Therefore, when non-duplicate epitopes were chosen, the reduction of agonistic activity correlated with the reduced size of the immunocomplexes. This result is consistent with the anti-TNFR2 BpAb we previously reported (21).

**Fig. 6.**
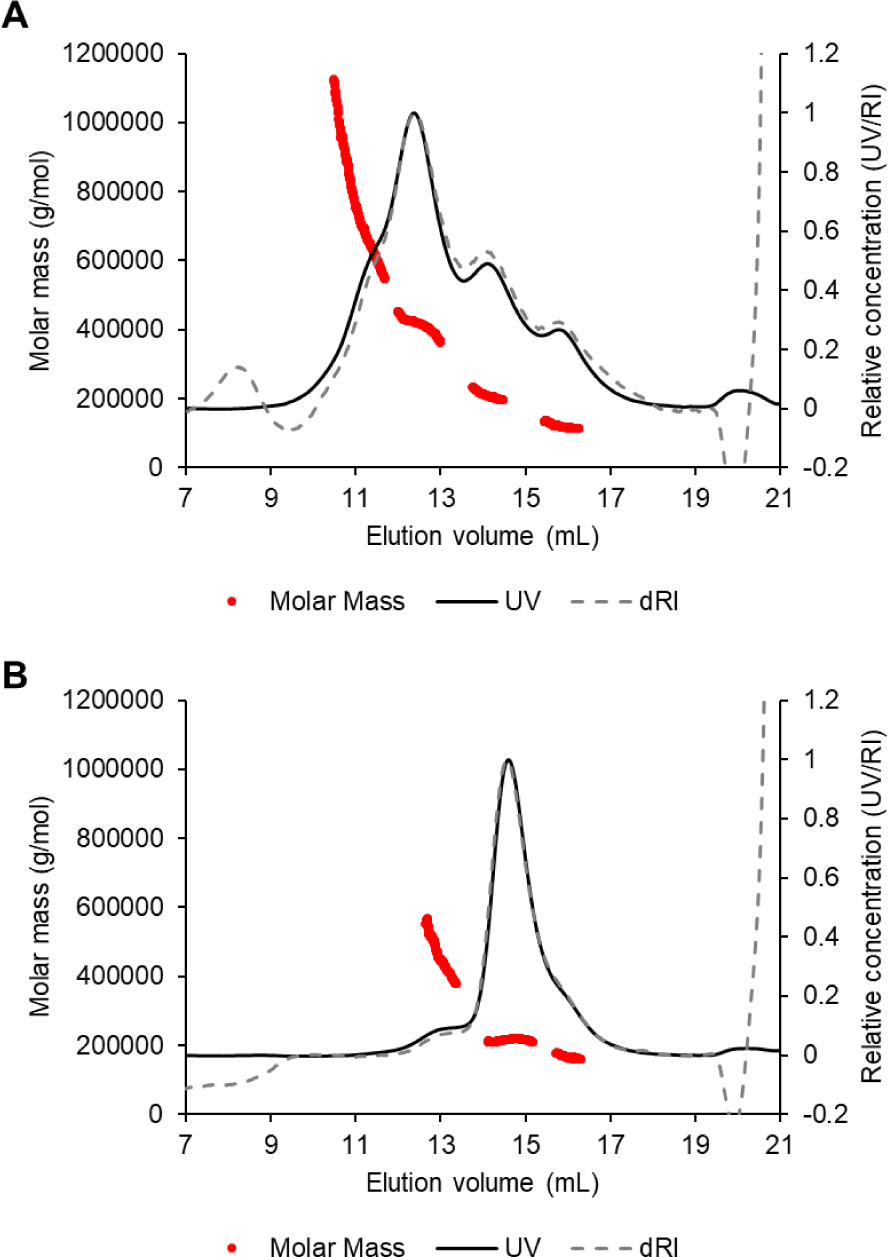
Charts of size-exclusion chromatography–multi-angle light scattering. A) BpT10-104, B) BpT10-405 in complex with CD30-MBP. See Supplementary Fig. S14 for each component.

## Discussion

The TNFRSF encompasses a broad and varied array of cell surface receptors that play pivotal roles in governing immune responses, inflammation, and cellular viability. Recent research has emphasized the significance of receptor clustering in both activating and modulating TNFRSF receptor signaling (11-13,33). Malfunction in TNFRSF signaling is linked to numerous pathological conditions (34). Thus, control of signaling has gathered attention for the development of therapeutics (11,12,34). Here, antibodies are promising (33); however, it has not been easy to design either agonists or antagonists logically. Although recent studies indicated strategies to maximize biological activities by structural optimization of conventional antibodies (35-38), it is still difficult to design them based on the mechanisms. Here, BpAbs offer attractive choice for the control (20,21,39).

For the logical development of BpAbs, understanding the mechanism of signal activation based on the relationship between structure and activity is crucial (19,20). As depicted in Fig. 1, the cluster-forming ability of cAbs and BpAbs affects signaling activity. Here, the stoichiometry of antibody-antigen complexes facilitates the initial level of understanding. Based on the epitope information, possible binding stoichiometry for all cAbs and BpAbs is categorized in Fig. 7A. cAbs are categorized into duplicate binders and others. Six cAbs binding non-duplicate epitopes have only one epitope per CD30 molecule, thus can form antibody:antigen = 1:2 complexes (Fig. 7B). Three cAbs binding the duplicate region have two epitope positions per CD30 molecule and can form n:n (n = 1,2,3,4…) complexes (Fig. 7C). For BpAbs, there are three categories, with zero, one, or two Fvs binding duplicate epitopes. Fifteen BpAbs using Fvs binding non-duplicate epitopes have two epitope positions and form immunocomplexes in the same manner as duplicate-binding cAbs (Fig. 7C). Three BpAbs using both Fvs from the duplicate region have four epitope positions per CD30 molecule, and they may exhibit the stoichiometry of antibody:antigen = 2n:n (n = 1,2,3,4…) (Fig. 7D). Finally, eighteen BpAbs using one Fv from the duplicate binder and another Fv from the non-duplicate binder have three epitope positions per CD30 molecule, and they may show complicated binding patterns with excess arms potentially binding the non-duplicate epitope (Fig. 7E).

**Fig. 7.**
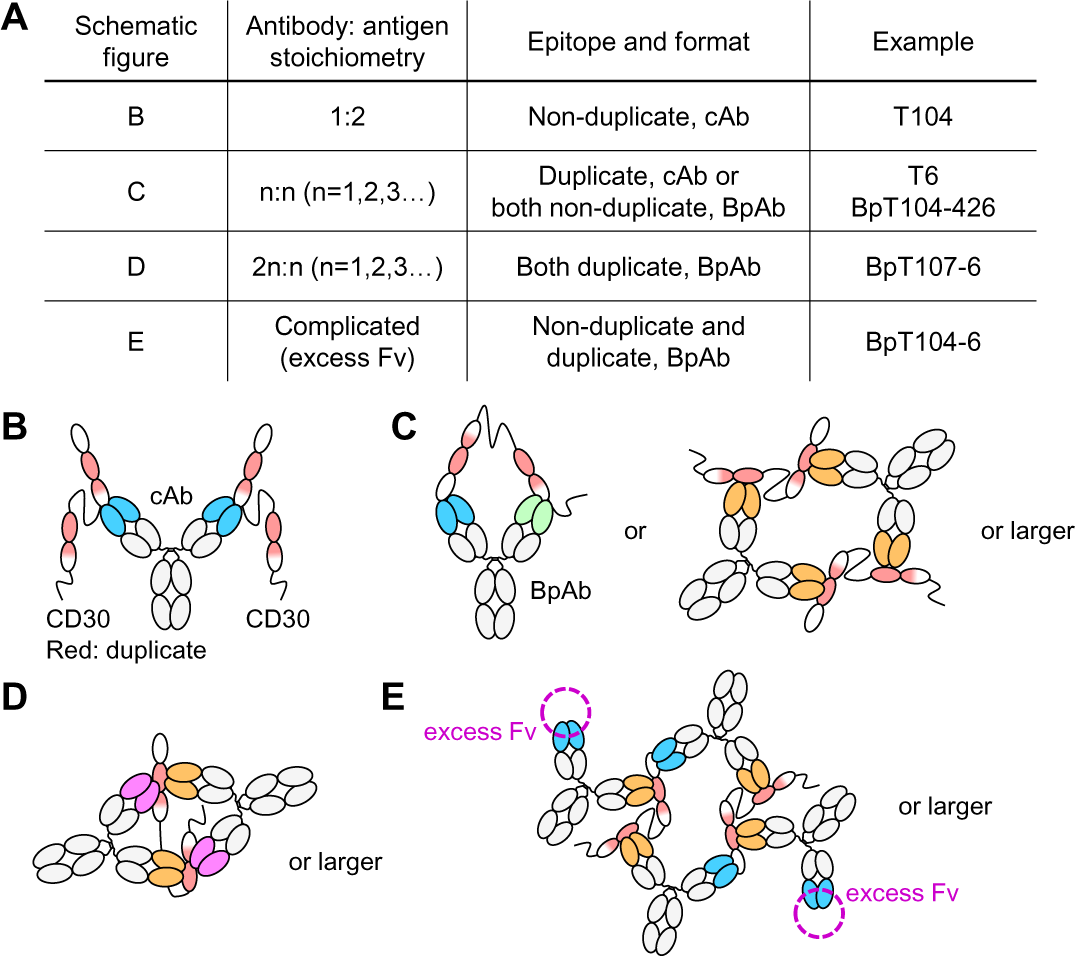
Possible binding stoichiometry by cAbs and BpAbs. A) Summarized table. B) cAbs recognizing non-duplicate epitopes show the stoichiometry of Ab:Ag = 1:2 (Ab, antibody; Ag, antigen). C) cAbs recognizing duplicate epitopes and BpAbs with two different Fvs each recognizing non-duplicate epitopes show the stoichiometry of Ab:Ag = 1:1. D) BpAbs with two different Fvs each recognizing duplicate epitopes show the stoichiometry of Ab:Ag = 2:1. E) BpAbs with two different Fvs each recognizing duplicate and non-duplicate epitopes do not show stable stoichiometry due to excess Fv recognizing non-duplicate epitopes.

As previously reported by us and others (21,39), the size of immunocomplexes plays a crucial role in determining the signaling activity of agonistic BpAbs against TNFRSF. Here, we initially focus on BpAbs that bind to non-duplicate epitopes (Fig. 7C). Among the six antibodies binding non-duplicate epitopes, four (T104, AC10, T426, and T105) bound to CRD1-3. For the six BpAbs produced from these four antibodies, a similar cluster-forming mechanism as observed for TNFR2 was expected. When viewed from the top of CRDs, the epitopes of AC10, T105, T426, and T104 are arranged in this order (Fig. 8A). We anticipated that the expected agonistic activity would be stronger for distal epitope pairs in this mapping. Indeed, the BpAb with the farthest pair of epitopes (AC10 and T104) exhibited the strongest signaling activity through cluster formation, as observed in SEC-MALS (Fig. 6A, Fig. 8B), suggesting a similar activation mechanism of TNFRSF by BpAbs (21,39). The other BpAbs demonstrated moderate agonistic activity, although it is challenging to discern slight differences among them. Due to the narrow non-duplicate region, Fv combinations such as T426 and T105 resulted in competition between each other (Tables 1,2); thus, the binding mode was similar to that of cAbs in these cases.

**Fig. 8.**
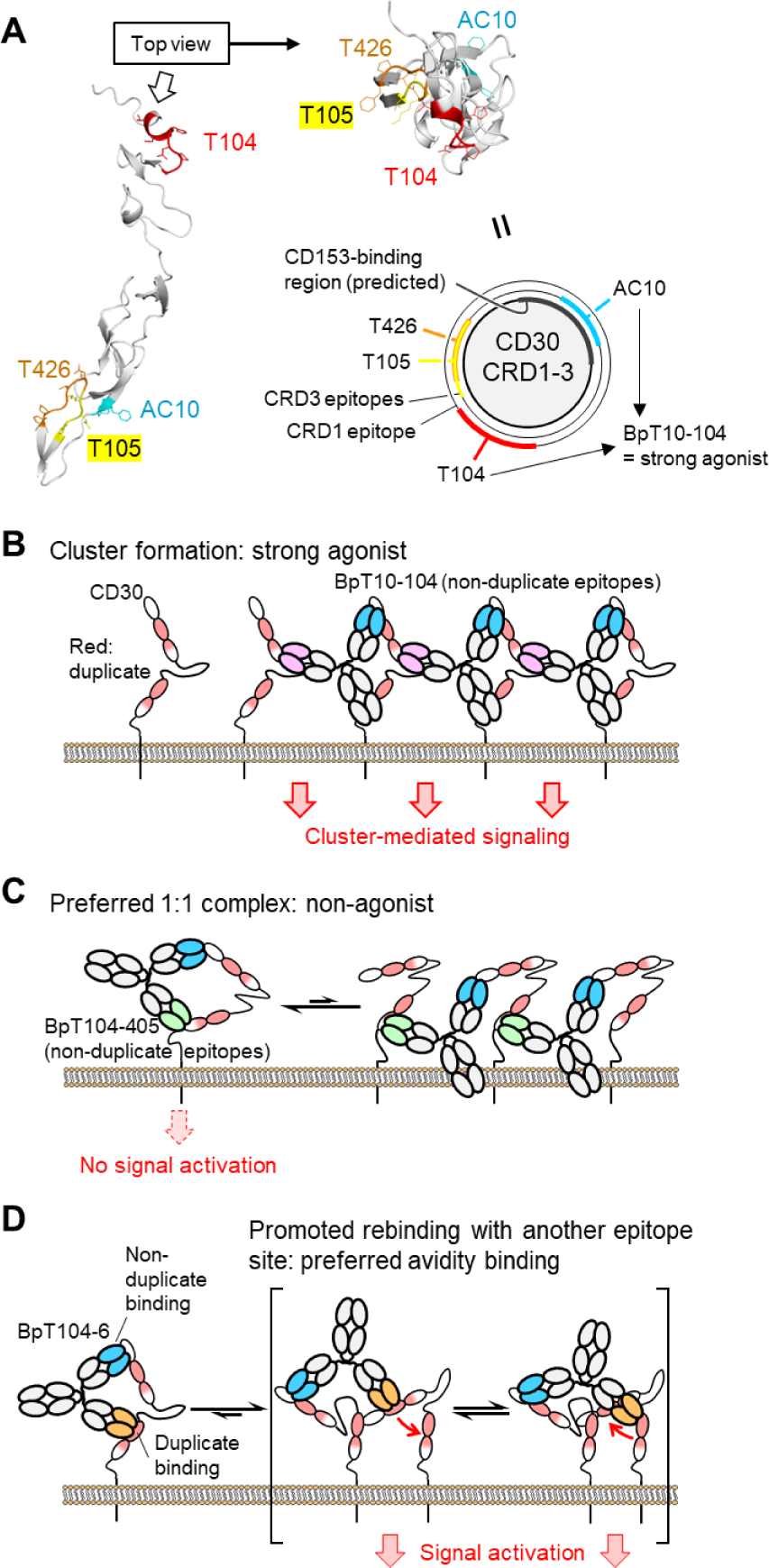
Proposed mechanisms of promoted and reduced agonistic activities among BpAbs. A) Relationship between the positions of four CRD1-3 non-duplicate epitopes and agonistic activities of BpAbs. B) BpT10-104, binding two regions in CRD1-3 with preferred cluster formation. C) Non-duplicate BpAbs divided by a flexible region which prefer 1:1 immunocomplex formation to show reduced agonistic activity. D) Putative mechanism of duplicate-binding BpAbs showing agonistic activity. Due to proximity, promotion of rebinding after dissociation of duplicate-binding Fv is expected. This may function as avidity binding with the preference of crosslinking multiple CD30 molecules over binding to a single CD30 molecule.

On the other hand, the other two non-duplicate antibodies (T25 and T405) bound to the epitopes in CRD6. Because of the flexible region between the two epitopes, the nine BpAbs with one of the Fvs from T25 or T405 were not strongly agonistic. In particular, BpT426-25, BpT426-405, BpT10-25, and BpT10-405 were non-agonists. Based on the analysis of BpT10-405 in SEC-MALS, they preferred the formation of 1:1 immunocomplexes (Fig. 6B, Fig. 8C). Since AC10 blocks CD153 binding (Fig. 5), BpT10-25 and BpT10-405 exhibited strong antagonistic activity, producing a pure antagonist in a similar manner as we previously reported for a BpAb against TNFR2 (21).

The 21 duplicate-binding BpAbs exhibited complicated activation patterns, likely due to simultaneous binding of three or four Fvs to a single CD30 molecule (Fig. 7D,E). The possible binding patterns are numerous, and generating a comprehensive model is challenging. There was no simple correlation with the relative position of epitopes in the activity. Interestingly, only two non-agonists (BpT10-107 and BpT107-427) were found among these BpAbs. We originally expected that BpAbs with at least one Fv binding the duplicate region may favor a 1:1-binding mode, thus most of these BpAbs may fall into non-agonists in a similar mechanism as shown in Fig. 8C. Experimental observation, in contrast with the prediction, suggested that most of these BpAbs would favor crosslinking between multiple CD30 molecules. The mechanism can be partly described by promoted rebinding after dissociation between two duplicate epitopes, resulting in avidity binding advantageous for a crosslinking binding pattern (Fig. 8D). Another observation is that only a small number of strong agonists were found, which would be consistent with this model due to the relatively low frequency of selecting a specific structure optimal for strong signal activation. These observations suggest difficulties in designing functional BpAbs targeting duplicate regions of CD30. The mechanisms may be shared with cAbs binding duplicate regions, which showed moderate agonistic activities.

Findings of the present study is similar to our investigation targeting TNFR2 (21). We discovered potential antagonists among 1:1-binding BpAbs, where two different Fvs bind different epitopes, which do not induce signal activation in the absence of a specific ligand. Such effective antagonists were not observed among cAbs. A significant revelation was the discovery of antagonistic activity in AC10-based BpAbs. AC10, the strongest agonistic cAb among analyzed, exhibited maximum signal activation surpassing even the strongest agonistic BpAb. However, by transitioning into a BpAb format, its potential to block CD153 was fully utilized by reduction of crosslinking. It is noteworthy that modification of the activity based on the flexible nature of CD30 is possible by using a BpAb format.

In summary, we developed 36 BpAbs from nine Fvs binding different epitopes of CD30, including three Fvs binding the duplicate region. We identified several BpAbs as non-agonists. Successfully, we developed strongly antagonistic BpAbs among CD153-competitive BpAbs. These BpAbs included those utilizing the Fv sequence from a potent agonist, AC10. The mechanism underlying reduced agonistic activity is akin to that observed in TNFR2, another member of TNFRSF, forming BpAb:antigen = 1:1 immunocomplex. These findings pave the way for the development of CD30 antagonists and also contribute to the design of antagonistic BpAbs against association-activated receptors.

## Materials and Methods

### Cell culture

HEK293T cells were sourced from the American Type Culture Collection and maintained in DMEM supplemented with 10% fetal bovine serum (FBS). KARPAS 299 cells were obtained from the German Collection of Microorganisms and Cell Cultures (DSMZ, Braunschweig, Germany). Ramos-Blue cells were purchased from InvivoGen (San Diego, CA, USA) and cultured in IMDM supplemented with 2 mM glutamine and 10% FBS. Ramos-blue cell line is an engineered human Ramos B cell (CD30-negative) containing an NF-B/AP-1-inducible secreted embryonic alkaline phosphatase (SEAP) gene. The cells were further transfected in our laboratory with pcDNA6-derived plasmids encoding full length of CD30. After transfection by electroporation (Nucleofector, Amaxa), a stable transformant was established by selection with blasticidin-containing medium and cloning via limiting dilution.

### Production of recombinant antibodies

Recombinant antibodies were produced according to established protocols (21,31). Conventional natural format of Abs (cAbs) were generated as human IgG1κ chimera antibodies in this study. Heavy and light chains were encoded in pFuse-CHIg-hG1 and pFUSE2-CLIg-hK vectors (InvivoGen), respectively. Recombinant cAbs were produced using the Expi293 Expression System (Thermo Fisher Scientific) and purified from the culture supernatant using rProtein A Sepharose Fast Flow column (Cytiva). Further purification was carried out using a HiLoad Superdex200 16/600 column. The expression and purification of BpAbs followed previously described a series of methods, which are based on intein-mediated protein trans-splicing (IMPTS) (21,31). Briefly, Fab ‘**N**’ was fused to Cfa Int^N^-MBP, and Fab ‘**C**’ was fused to an Fc with a ‘knob’ mutation using the knobs-into-holes method. Fab ‘**C’** was co-expressed with an MBP-Cfa Int^C^ fusion of Fc with a ‘hole’ mutation to produce a monovalent antibody. Following IMPTS, the BpAbs were purified using a Superdex 200 Increase 10/300 GL column (Cytiva).

### Preparation of recombinant CD30 and CD153 proteins

CD30 whole extracellular domains (NP_001234.3, amino acids 1-383), or CD30 fragments (CRD4, CRD1-3, or CRD4-6) with Ig-κ chain signal peptide for the secretion at the fronts were genetically fused to a peptide linker SGRGGRRASVPDPEG followed by the hinge and Fc portion of human or rabbit IgG1, and cloned into EcoRI-XbaI site of pcDNA3 plasmid (InVitrogen). The purified plasmids were transfected into 293T cells and the fusion proteins were harvested from the culture supernatant (40). The secreted CD30-rFcs contained DAAQP at the N-terminus after signal peptide cleavage.

CD30 extracellular domains (1-336) fused to MBP as a tag (CD30-MBP; Supplementary Fig. S1) were cloned into pcDNA3.1. The protein was expressed in Expi293 Expression System. The culture supernatant was dialyzed overnight against Buffer A (20 mM Tris, 300 mM NaCl, pH 8.0) containing 5 mM imidazole. Expressed proteins were captured on Ni-NTA Superflow (Qiagen) and subsequently washed with Buffer A containing 5, 10, and 20 mM imidazole, and the protein was eluted using Buffer A containing 200 mM imidazole. The eluate was dialyzed against phosphate-buffered saline (PBS) and the final purification was conducted using a Superdex200 16/600 column.

CD30 ligand (CD153) is a type II membrane protein whose C-terminus is exposed to the outside of cells. The cDNA coding Fc portion of rabbit IgG (P01870.1 amino acid 96-323,was fused to the N-terminus of cDNA coding the extracellular domain of human CD153 (NP_001235.1, Gln63-Asp234), and subcloned into SfiI-XhoI site of pSecTag2 Hygro A plasmid (InVitrogen). As a result, the secreted CD153-rFc contains DAAQP at the N-terminus upon cleavage of the signal peptide, the rabbit IgG Fc, GGPRSDLEVLFQGPLGSGG as a linker, and 63-234 of CD153. The Fc-fused proteins were expressed in HEK293T cells and the fusion protein in the culture supernatant was purified by protein A affinity chromatography as described previously (22).

## Mutual competitive binding assay of cAbs

The nine anti-CD30 cAbs were selected based on the topographical epitopes previously identified by various conditions of competition assays (24-26). To reevaluate the topographical relationships of the epitopes, a competitive ELISA was performed under a single condition, including all nine cAbs in this study. All pairs of the 9 cAbs (9x9) were tested with the sequential binding activity for the binding to CD30-rFc protein as described previously with slight modifications (21,24). Microplates (MaxiSorp, Nalge Nunc, Rochester, NY) were coated with goat anti-human IgG Fc (#109-005-098, Jackson Immunoresearch) and each indicator cAb (0.75 μg/mL) was captured. Competitor cAb (2.8 μg/mL) and CD30-rFc (50 ng/mL) were incubated overnight at 4 °C in a separate tube. The plates were washed twice and the competitor–CD30-rFc solution was added to each well of the coated plates. After washed twice, bound CD30-rFc was probed by alkaline phosphatase-conjugated goat anti-rabbit IgG (#111-055-046, Jackson Immunoresearch).

For SPR-based analyses, F(ab′)_2_ protein was produced by digestion using 10% (w/w) IdeS protease on each cAb. His-tagged IdeS protease and digested Fc were removed by NEBExpress Ni Spin Columns (New England Biolabs) and Protein A HP SpinTrap (Cytiva), respectively. cAbs and CD30-MBP were successively captured as described above, and the interaction of F(ab′)_2_ (20 nM) was analyzed by the contact and dissociation time of 90 and 120 s, respectively, flowed at 30 μL/min.

### Binding analysis of CD153 to CD30

The reactivity of CD153-rFc to CD30 or CRDs fused to human Fc were tested in sandwich ELISA. Microplates were coated with goat anti-human IgG Fc antibody (#109-005-098, Jackson Immunoresearch) for 2 h at room temperature. After blocking (25% DMEM, 5% FBS, 0.5% bovine serum albumin, and 0.1% sodium azide in PBS) and washing, to appropriate wells we added 200 ng/100 µL/well of each CD30-deletion mutant-human Fc in blocking buffer. After washing, CD153-rFc in concentration series was added. HRP-goat-anti-rabbit IgG Fc antibody (#111-035-046, Jackson Immunoresearch) was used to detect the bound CD153-rFc.

### CD30 orthologs

Ortholog species containing the *tnfrsf8* gene with moderate structural homology to human CD30 (Refseq NP_001234.3) were screened. Homologous sequences within the extracellular region of human CD30 (F19-G385 as defined in UniProt) were identified using protein BLAST (30). Seven sequences with approximately 60% homology to human CD30 (Refseq XP_022454406.1, XP_036855465.1, XP_014705247.1, KAF6380682.1, VFV35557.1, XP_006197082.2, XP_013359778.1) and five sequences with approximately 50% homology to human CD30 (Refseq XP_036084707.1, XP_027474377.1, XP_007533056.1, XP_034527286.1, ELW68632.1) were selected. Among the 12 sequences, five with distinct distances from each other in the phylogenetic tree generated using Clustal 2.1 were further chosen for mapping antibody binding (41). DNA encoding the full-length TNFRSF8 proteins of these five orthologs (KAF6380682.1, VFV35557.1, XP_006197082.2, XP_013359778.1, ELW68632.1) was synthesized by Genewiz LLC.

### Structure prediction

The predicted human CD30 structure was obtained from AlphaFold DB (AF-P28908-F1) (28). Structures of CD30 orthologs from other species were predicted using AlphaFold2 in ColabFold v1.1 with the MSA mode of MMSeq2 (27,42). One of the five generated models was visualized using Pymol (Schrödinger, LLC).

### Ortholog and epitope mapping

Epitopes of the antibodies were determined following a similar protocol as described previously (21). DNA encoding human and the five ortholog sequences were fused with TagBFP on the C-terminus to ensure the expression of folded proteins on the cell membrane, and then cloned into pcDNA3.1. Mutants with partial substitution with the ortholog sequences were also generated either by seamless cloning using the In-fusion HD Cloning Kit (Takara) or by site-directed mutagenesis using the KOD-Plus Mutagenesis Kit (Toyobo). The sequences of the encoded proteins used in the ortholog and epitope mapping are described in Table S1. The expression vector was transfected into HEK293T cells using PEI ‘MAX’ (Polysciences Inc.). Cells were cultured for 40 h post-transfection and detached from the culture vessels using trypsin/EDTA for 5 min. Cells were covalently labeled with combinations of succinimidyl ester compounds of Pacific Orange, DyLight 633, or DyLight 800 (Cat. No. P30253, 46414, and 46421; Thermo Fisher Scientific) as described previously (21). Twelve differently labeled cells at maximum per experiment were combined, and the cells were then incubated with the cAbs (1.5 μg/mL) and the secondary antibody anti-human IgG, Fcγ-PE (1/200 dilution, #109-116-170; Jackson Immunoresearch). The cells were analyzed using a BD LSRFortessa Cell Analyzer (BD Biosciences), and the acquired data were analyzed using FlowJo_v10.9.0 (FlowJo, LLC). Antibody binding as a function of PE fluorescence relative to the expression of CD30-TagBFP was analyzed. All antibodies were always analyzed at the same time because antibodies unaffected by mutation can function as the positive control.

### Reporter assay

CD30-expressing Ramos-Blue (CD30-RB) cells were seeded at 5 × 10^4^ in 100 μL of medium-containing antibodies at the indicated concentrations (100 fM to 100 nM in tenfold dilution series) in the absence or presence of 200 ng/mL CD153-rFc. The cells were incubated for 18 h, and secreted alkaline phosphatase was analyzed using *p*-nitrophenyl phosphate. Colorimetric changes were determined by measuring absorbance at 405 nm using an EnSpire microplate reader (PerkinElmer). Signals were normalized to the average absorbance of the eight wells incubated without antibody or CD153-rFc (negative) and the eight wells incubated only with 50 ng/mL CD153-rFc (positive). For the chart showing the agonistic and antagonistic activities, the negative and positive values were set at 0.1 and 0.9, respectively.

### Flow cytometry

For antibody binding, KARPAS 299 cells or CD30-RB cells (1 × 10^5^ cells/well) were incubated with each antibody for 30 min on ice. After being washed twice, the cells were further incubated with the secondary antibody anti-human IgG, Fcγ-PE (1/200 dilution, #109-116-170; Jackson Immunoresearch) for 30 min on ice. For CD153-binding competition, CD30-RB cells were incubated with a mixture of the antibodies (1.5 μM) and CD153-rFc (200 ng/mL) for 30 min on ice, washed twice, and further incubated with the secondary antibody anti-rabbit IgG-PE (1/200, #111-116-144; Jackson Immunoresearch). The cells were then analyzed using a BD LSRFortessa Cell Analyzer. Data were analyzed in FlowJo v10.10.0.

### Binding affinity analysis to recombinant CD30 protein in SPR

The binding kinetics of an antibody-antigen interaction was measured using a Biacore T200 instrument (Cytiva). In brief, anti-human Fc was immobilized on a CM5 chip using the Human Antibody Capture Kit (Cytiva). To immobilize anti-human Fc (approximately 7000 RU), cAbs and BpAbs (1 μg/mL) were capture by flowing at 10 μL/min for 300 s (T25 and BpT104-405) or 60 s (others). The interaction with CD30-MBP was analyzed by the contact and dissociation time of 90 and 300 s, respectively, flowed at 30 μL/min. The concentration of CD30-MBP was 0, 20, and 200 nM for analyzing T426 and 0, 2, and 20 nM for analyzing others.

### Size-exclusion chromatography with multi-angle light scattering detector (SEC-MALS)

SEC-MALS measurements were conducted as previously described (21). Briefly, each antibody and CD30-MBP were mixed equimolarly (2 μM) in PBS, and a 50 μL solution was loaded onto a Superose 6 Increase 10/300 GL column (Cytiva) column. Light scattering was detected in DAWN 6 (Wyatt, Santa Barbara, CA, USA). Data were analyzed using ASTRA 6 software (Wyatt). The protein concentration was calculated from the refractive index using dn/dc = 0.175. Molar mass values were determined by the Debye fitting of angle-dependent light scattering.

## Acknowledgments

This work was partly supported by AMED grant numbers JP22ak0101099 and JP24ama121042, JSPS grant number JP24K01270, Takeda Science Foundation, and Kyoto University Foundation.

## Author contribution

Hiroki Akiba (Conceptualization, Funding Acquisition, Investigation, Writing-Original Draft), Tomoko Ise (Investigation, Writing-Review & Editing), Reiko Satoh (Investigation), Yasuhiro Abe (Investigation), Kouhei Tsumoto (Methodology), Hiroaki Ohno (Writing-Review & Editing), Haruhiko Kamada (Investigation, Funding Acquisition, Writing-Review & Editing), and Satoshi Nagata (Conceptualization, Methodology, Writing-Original Draft).

## Data Availability

All data supporting the findings of this study are available within the article and Supplemental Information. Raw data files are available by reasonable request to the corresponding authors.

## Ethics and Consent Statement

Not applicable.

## Animal Research Statement

Not applicable.

