## Supplementary Materials for "Generation of antagonistic biparatopic anti-CD30 antibody from an agonistic antibody by precise epitope determination and utilization of structural characteristics of CD30 molecule"

| Table of Contents | Page # |
|---|---|
| Supplemental Text | S2-3 |
| Table S1–S3* | S4-5 |
| Fig. S1–S14 | S6-21 |

* Table S1 is not included and provided as a separate file.

**Information of epitopes of anti-CD30 antibody clones unused in this study**

In the present study, a panel of 9 MAbs to 9 distinct topographical epitopes on the conformational structure of CD30 was selected from previously known 42 mAbs. The selection process was informed by information obtained for the 42 known mAbs and conducted as eliminating redundant mAbs to the same epitopes. The epitope locations of the mAbs were defined relatively with those of the other anti-CD30 mAbs by competition assays using CD30-expressing cells or CD30-Fc fusion protein in the following previous reports.

Twelve reference mAbs have been assigned to human CD30 in previous Human Leucocyte Differentiation Antigens (HLDA) workshops (1). Out of the 12, AC10 clone was used in the final panel of 9 mAbs in this study; Ki-I (2), Ber-H2 (3) and HRS-4 (4) had been directly evaluated for the relative epitope location with the selected 9 mAbs by us using 28 different mAbs (5-7); HRS-1 (8) and HRS-3 were reported to bind to the same epitope group for Ber-H2 and HRS-4 (4); Ber-H8 had been known to recognize the same epitope for Ber-H2 (9); Ber-H6 and Ber-H10 bind to 1-93 amino acid region as Ki-I (10); BerH4 and Ber-H8 react with 112-412 amino acid resign (10). HeFi-I clone (11) is one of the initially isolated anti-CD30 mAbs and was used in a past phase I clinical trial (ClinicalTrials.gov Identifier: NCT00048880). HeFi-I has been often used as a reference antibody in previous studies to evaluate the topographical locations of other CD30 epitopes. HeFi-I was included in the final 18 mAbs panel in this study. HeFi-I binding for CD30 was reported to be inhibited by AC10 (12). Clones M44 and M67 (13) and clones Ki-2, Ki-3, Ki-4, Ki-5, Ki-6 and Ki-7 (12) were produced separately. A comprehensive epitope mapping was reported in (12), which indicates: (i) Ki-2, Ki-4, Ki-6 and Ki-7 recognize the same topographical epitope as Ber-H2, HRS-1 and HRS-4; (ii) M67 and Ki-5 bind to an epitope group defined by Ki-I; (iii) M44 and Ki-3 bind to the same epitope as AC10 and HeFi-I. A part of this epitope mapping results was supported by another study (9). The two epitopes defined for M67 and Ki-4 were included in the 9 topographical epitope groups based on our experiments (5-7). Another human anti-CD30 mAb, 5F11, was generated in the HuMAb mouse (14), and was used in clinical trials (15). 5F11 bind to the topographical epitope defined by Ber-H2, Ki-4, Ki-2 and HRS-3 but the location is distinct from the epitopes of Ki-1or of AC10 and Ki-3 (14). We have produced 23 anti-CD30 mAbs in 6 experiments (6,7).

**Table S1.** Sequences of the orthologs and mutants used for epitope determination (separate file)

**Table S2.** Production yields of $Int^N$- or $Int^C$-fused antibody fragments

| | mg/L culture |
|---|---|
| T104N | 52 |
| T426N | 86 |
| AC10N | 172 |
| T105N | 105 |
| T25N | 40 |
| T405N | 21 |
| T6N | 6 |
| T427N | 32 |
| T107N | 107 |
| T104C | 328 |
| T426C | 366 |
| AC10C | 305 |
| T105C | 397 |
| T25C | 165 |
| T405C | 393 |
| T6C | 297 |
| T427C | 242 |
| T107C | 367 |

**Table S3.** Kinetic parameters of antibody interaction with CD30 determined by surface plasmon resonance in a rough-level analysis

| Name | $k_{on}$ ($\times 10^6$ $M^{-1}s^{-1}$) | $k_{off}$ ($\times 10^{-3}$ $s^{-1}$) | $K_D$ (nM) | Name | $k_{on}$ ($\times 10^6$ $M^{-1}s^{-1}$) | $k_{off}$ ($\times 10^{-3}$ $s^{-1}$) | $K_D$ (nM) |
|---|---|---|---|---|---|---|---|
| T104 | 0.32 | 2.8 | 8.8 | BpT25-405 | 0.29 | 0.26 | 0.91 |
| T426 | N.D. | N.D. | N.D. | BpT107-6 | 0.73 | 0.27 | 0.37 |
| AC10 | 1.2 | 1.6 | 1.3 | BpT107-427 | 1.0 | 0.61 | 0.60 |
| T105 | 0.70 | 1.5 | 2.2 | BpT427-6 | 1.1 | 0.29 | 0.26 |
| T25 | 0.17 | 1.3 | 7.5 | BpT104-6 | 1.0 | 0.61 | 0.59 |
| T405 | 0.18 | 2.4 | 13 | BpT104-427 | 1.1 | 0.36 | 0.32 |
| T6 | 1.6 | 1.5 | 0.94 | BpT107-104 | 1.2 | 0.47 | 0.40 |
| T427 | 1.2 | 0.64 | 0.55 | BpT10-6 | 0.98 | 0.26 | 0.27 |
| T107 | 0.10 | 0.61 | 6.0 | BpT10-427 | 1.1 | 0.63 | 0.58 |
| BpT104-426 | 1.3 | 3.9 | 2.9 | BpT10-107 | 1.1 | 0.69 | 0.61 |
| BpT104-25 | 3.1 | 1.6 | 0.50 | BpT107-25 | 0.23 | 0.20 | 0.87 |
| BpT104-405 | 0.16 | 7.6 | 48 | BpT107-405 | 0.18 | 0.33 | 1.8 |
| BpT426-25 | 0.37 | 0.31 | 0.82 | BpT426-6 | 1.1 | 0.50 | 0.45 |
| BpT426-405 | 0.30 | 0.61 | 2.1 | BpT426-427 | 1.2 | 0.53 | 0.44 |
| BpT10-104 | 1.0 | 0.26 | 0.26 | BpT25-6 | 0.83 | 0.39 | 0.46 |
| BpT10-426 | 1.1 | 0.25 | 0.23 | BpT25-427 | 0.91 | 0.52 | 0.57 |
| BpT10-105 | 0.98 | 0.065 | 0.066 | BpT105-6 | 1.1 | 0.39 | 0.37 |
| BpT10-25 | 0.92 | 0.58 | 0.64 | BpT105-427 | 0.94 | 0.23 | 0.24 |
| BpT10-405 | 0.98 | 0.62 | 0.63 | BpT105-107 | 1.1 | 0.30 | 0.26 |
| BpT105-104 | 0.75 | 0.24 | 0.32 | BpT405-6 | 1.1 | 0.49 | 0.44 |
| BpT105-426 | 0.49 | 1.2 | 2.3 | BpT427-405 | 1.0 | 0.32 | 0.31 |
| BpT105-405 | 1.4 | 0.94 | 0.67 | BpT107-426 | 0.65 | 0.22 | 0.34 |
| BpT25-105 | 1.2 | 0.65 | 0.55 | | | | |

A

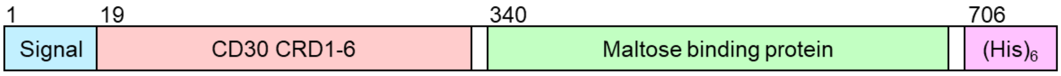


MRVLLAALGLLFLGALRAFPQDRPFEDTCHGNPSHYYDKAVRRCCYRCPMGLFPTQQCPQRPTDCRKQCEPDYYLDEADRC
TACVTCSRDDLVEKTPCAWNSSRVCECRPGMFCSTSAVNSCARCFFHSVCPAGMIVKFPGTAQKNTVCEPASPGVSPACAS
PENCKEPSSGTIPQAKPTPVSPATSSASTMPVRGGTRLAQEAASKLTRAPDSPSSVGRPSSDPGLSPTQPCPEGSGDCRKQ
CEPDYYLDEAGRCTACVSCSRDDLVEKTPCAWNSSRTCECRPGMICATSATNSCARCVPYPICAAETVTKPQDMAEKDTTF
EAPPLGTQPDCNGGSKIEEGKLVIWINGDKGYNGLAEVGKKFEKDTGIKVTVEHPDKLEEKFPQVAATGDGPDIIFWAHDR
FGGYAQSGLLAEITPDKAFQDKLYPFTWDAVRYNGKLIAYPIAVEALSLIYNKDLLPNPPKTWEEIPALDKELKAKGKSAL
MFNLQEPYFTWPLIAADGGYAFKYENGKYDIKDVGVDNAGAKAGLTFLVDLIKNKHMNADTDYSIAEAAFNKGETAMTING
PWAWSNIDTSKVNYGVTVLPTFKGQPSKPFVGVLSAGINAASPNKELAKEFLENYLLTDEGLEAVNKDKPLGAVALKSYEE
ELVKDPRIAATMENAQKGEIMPNIPQMSAFWYAVRTAVINAASGRQTVDEALKDAQGHHHHHH

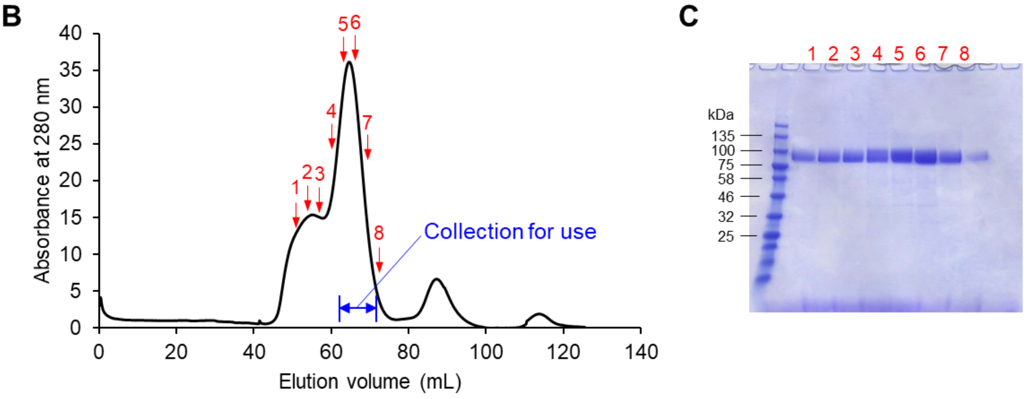


**Fig. S1.** Maltose-binding protein-fused CD30 (CD30-MBP). A) Construction of CD30-MBP and the peptide sequence. B) Size-exclusion chromatogram for the final step of CD30-MBP purification. C) SDS-PAGE analysis of peak fractions from size-exclusion chromatography. Red-numbered arrows in B correspond to lanes in C.

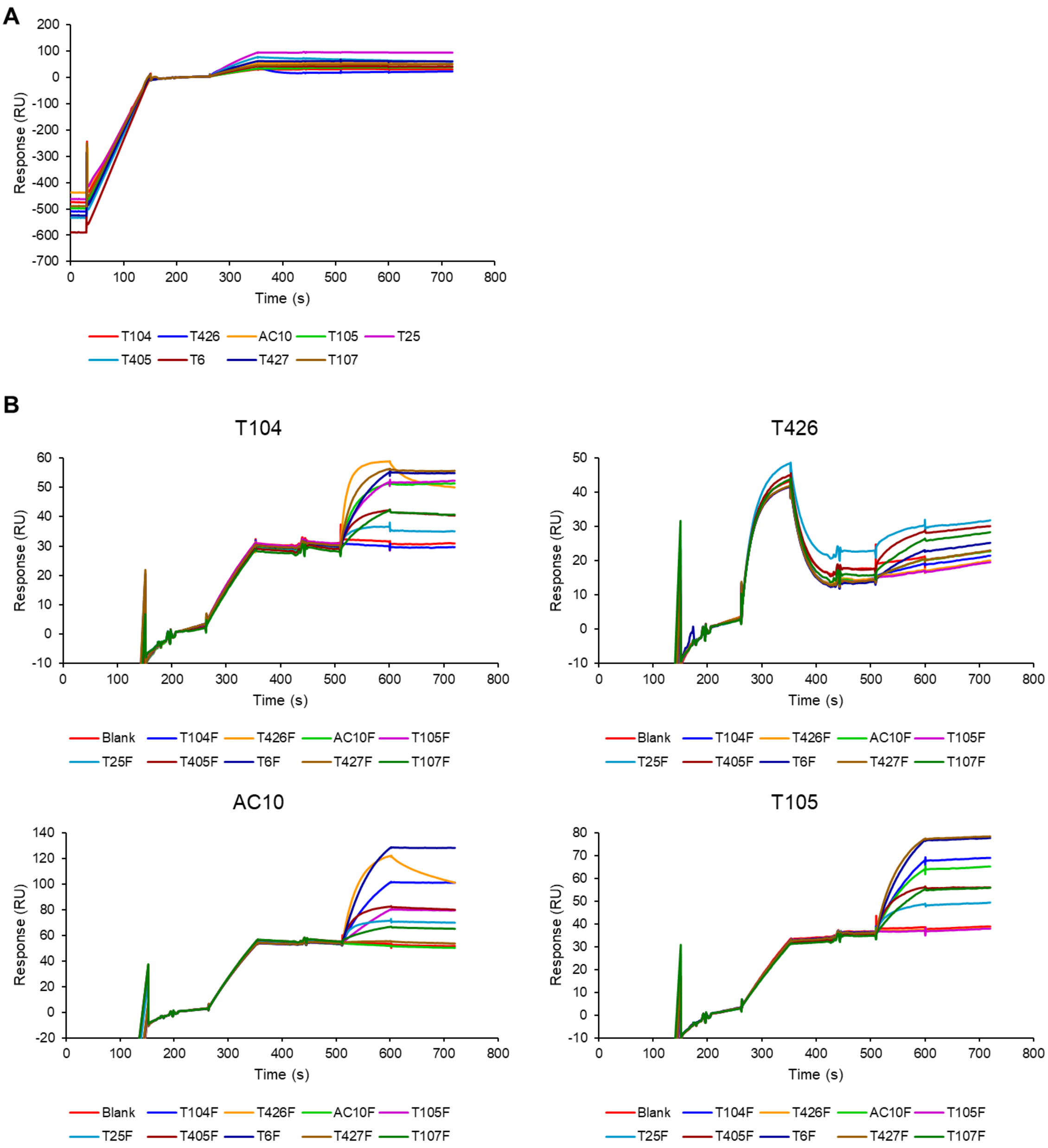
A
Response (RU)
Time (s)
T104
T426
AC10
T105
T25
T405
T6
T427
T107
B
T104
T426
AC10
T105
Blank
T104F
T426F
AC10F
T105F
T25F
T405F
T6F
T427F
T107F

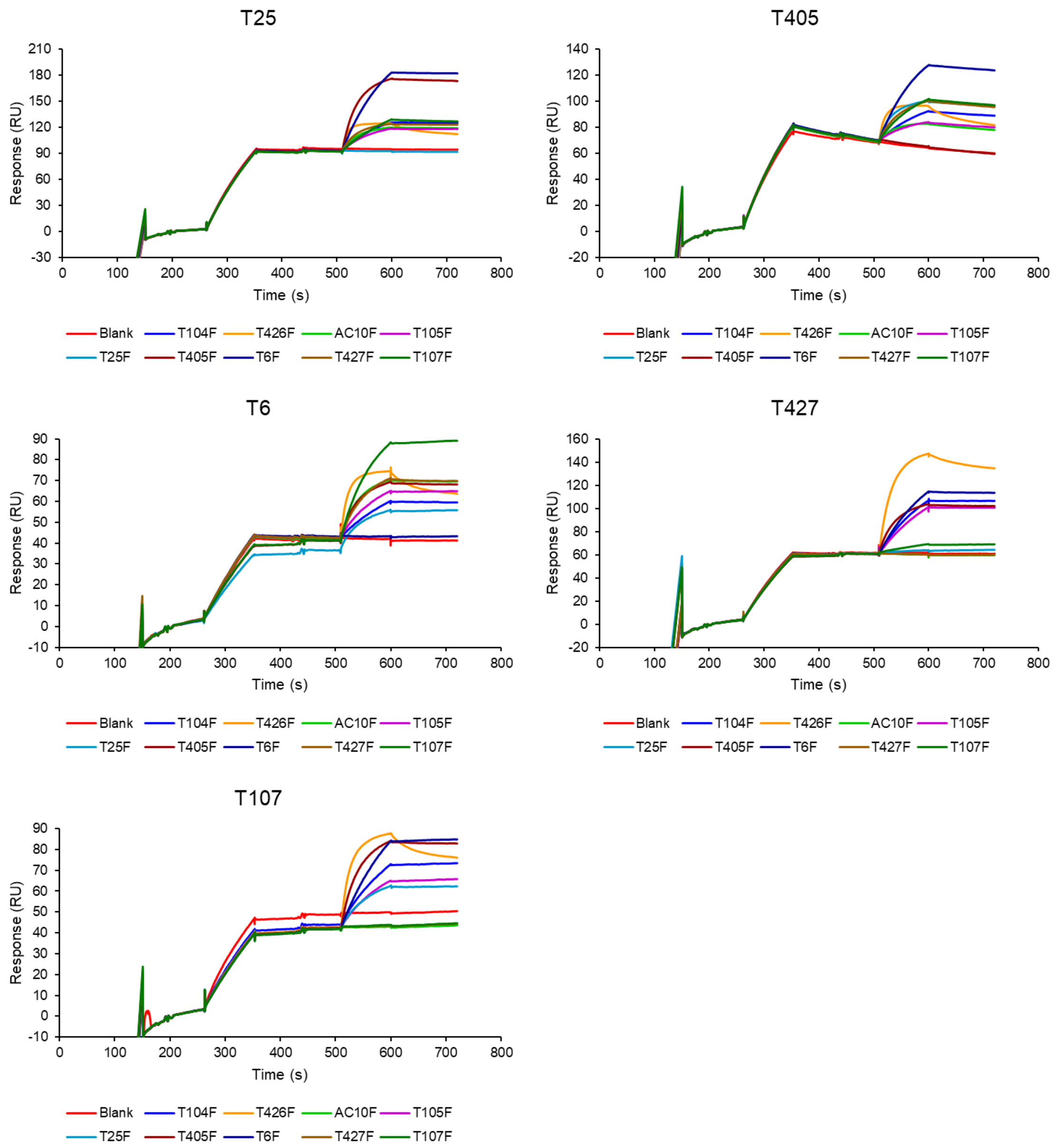


**Fig. S2.** Surface plasmon resonance sensorgrams for evaluating competitive binding by variable regions. A) Capture of cAbs by anti-Fc (30–150 s), followed by capture of CD30-MBP by the cAbs (270–350 s). These runs correspond to the 'Blank' runs in B. B) Interaction of F(ab′)$_2$ antibodies with CD30-MBP captured by the indicated cAbs on top of the panels. To capture CD30-MBP as described above, F(ab′)$_2$ antibodies were flowed to contact (510–600 s), followed by dissociation.

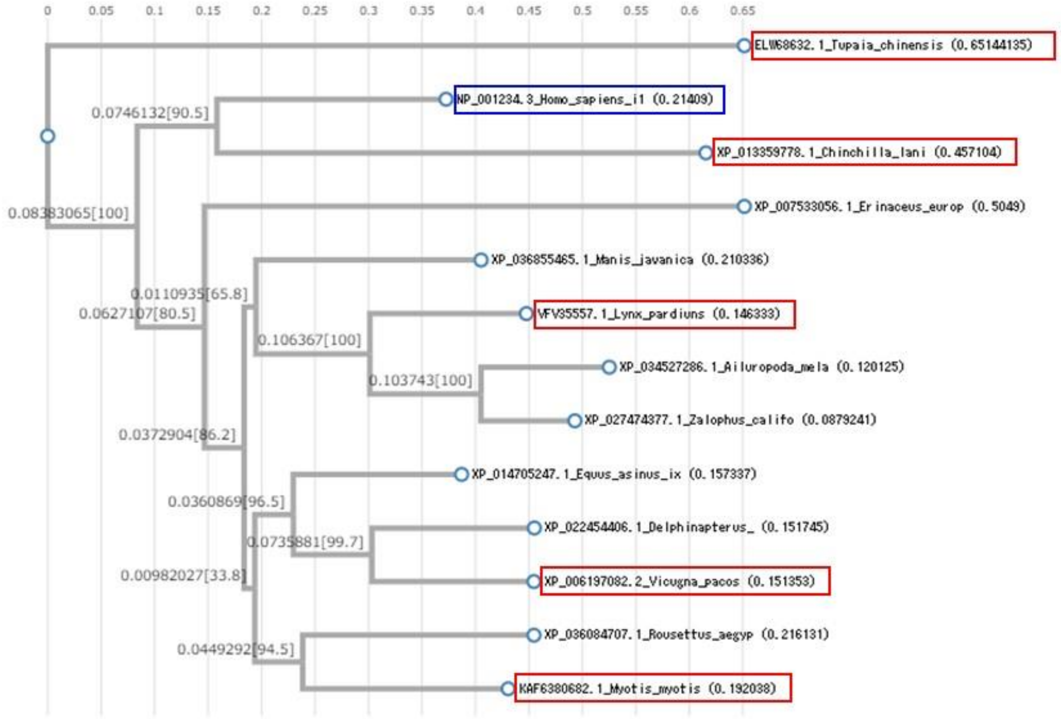


**Fig. S3.** Phylogenetic tree of human CD30 proteins (blue) and five orthologs (red) among selected sequences.

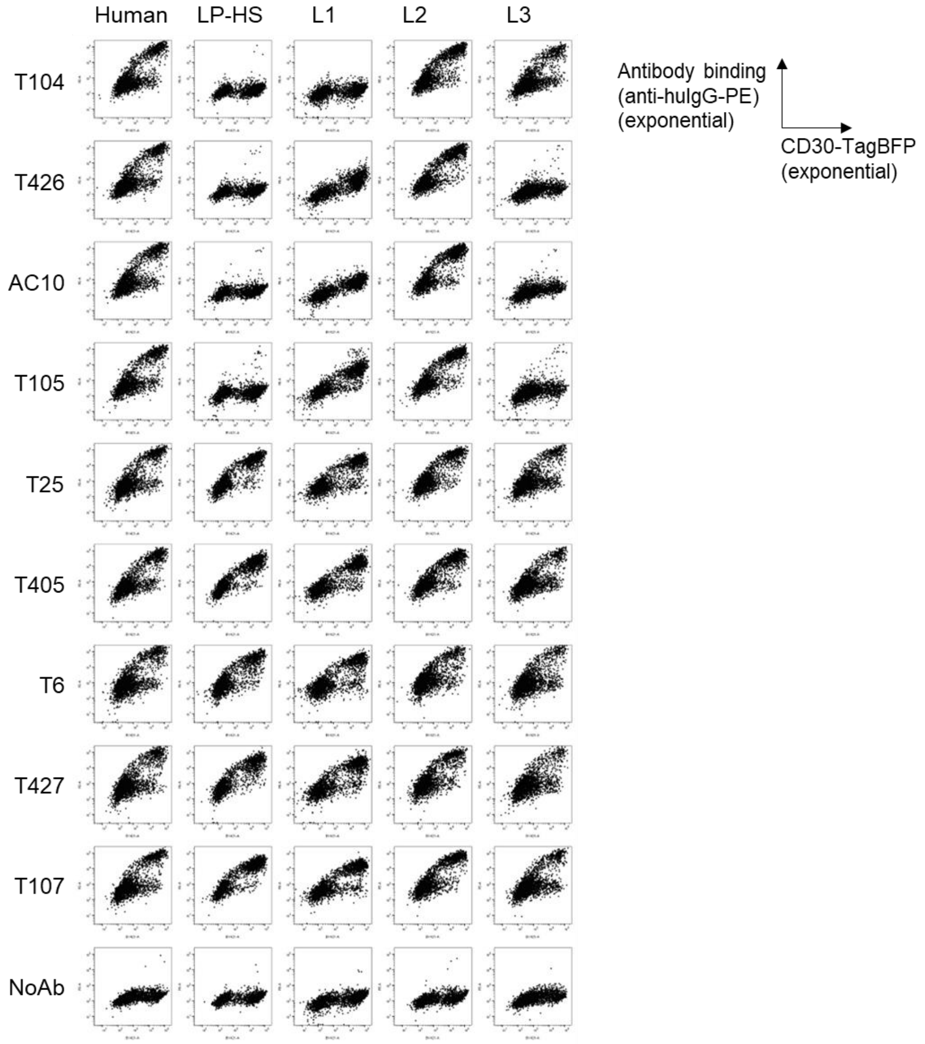


**Fig. S4.** Antibody binding to human CD30 and domain-substituted CD30 proteins with L.pard sequences in CRD1-3 (LP-HS), CRD1 (L1), CRD2 (L2), and CRD3 (L3).

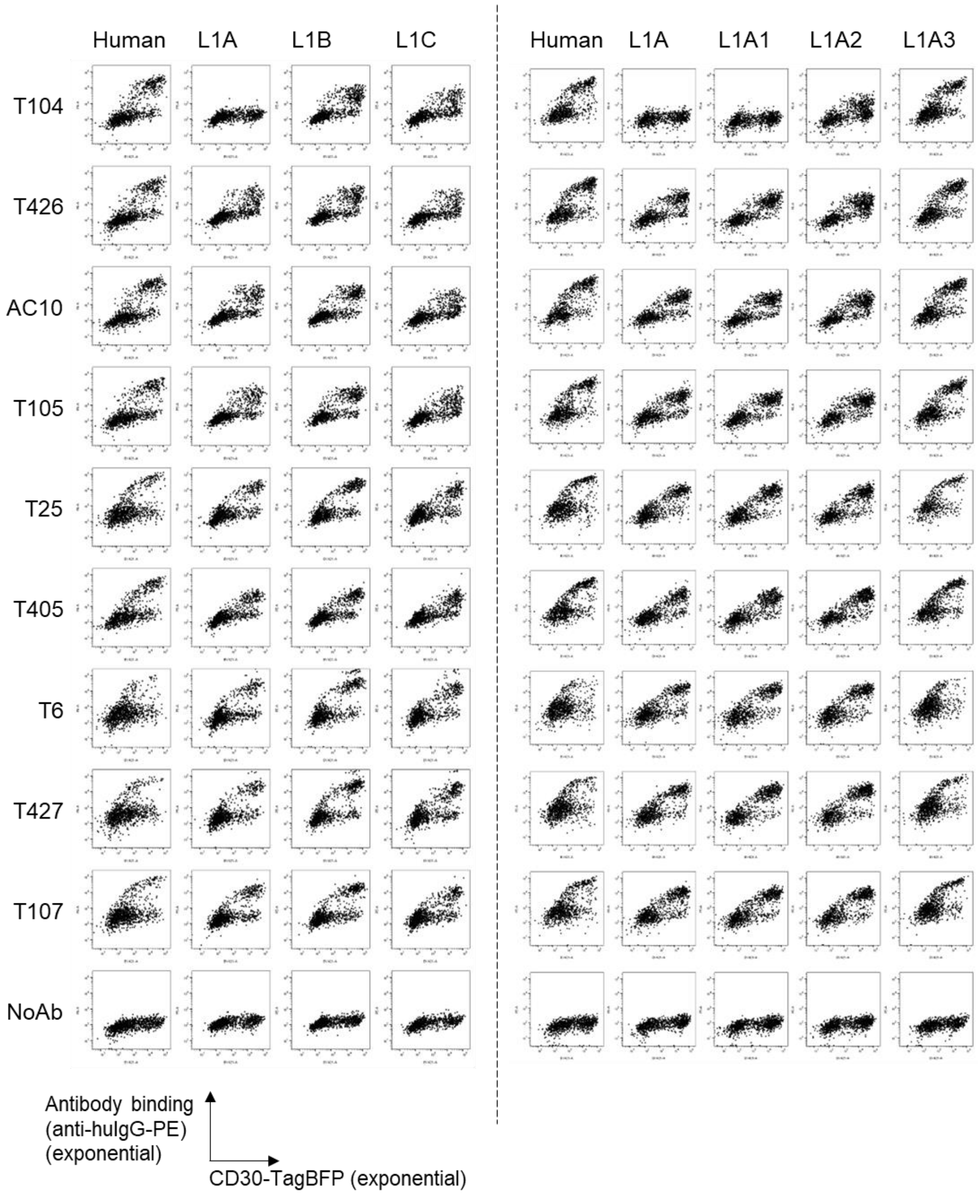


**Fig. S5.** Antibody binding to human CD30 with a portion of CRD1 substituted with L.pard sequences. Split line indicates data from independent experiments.

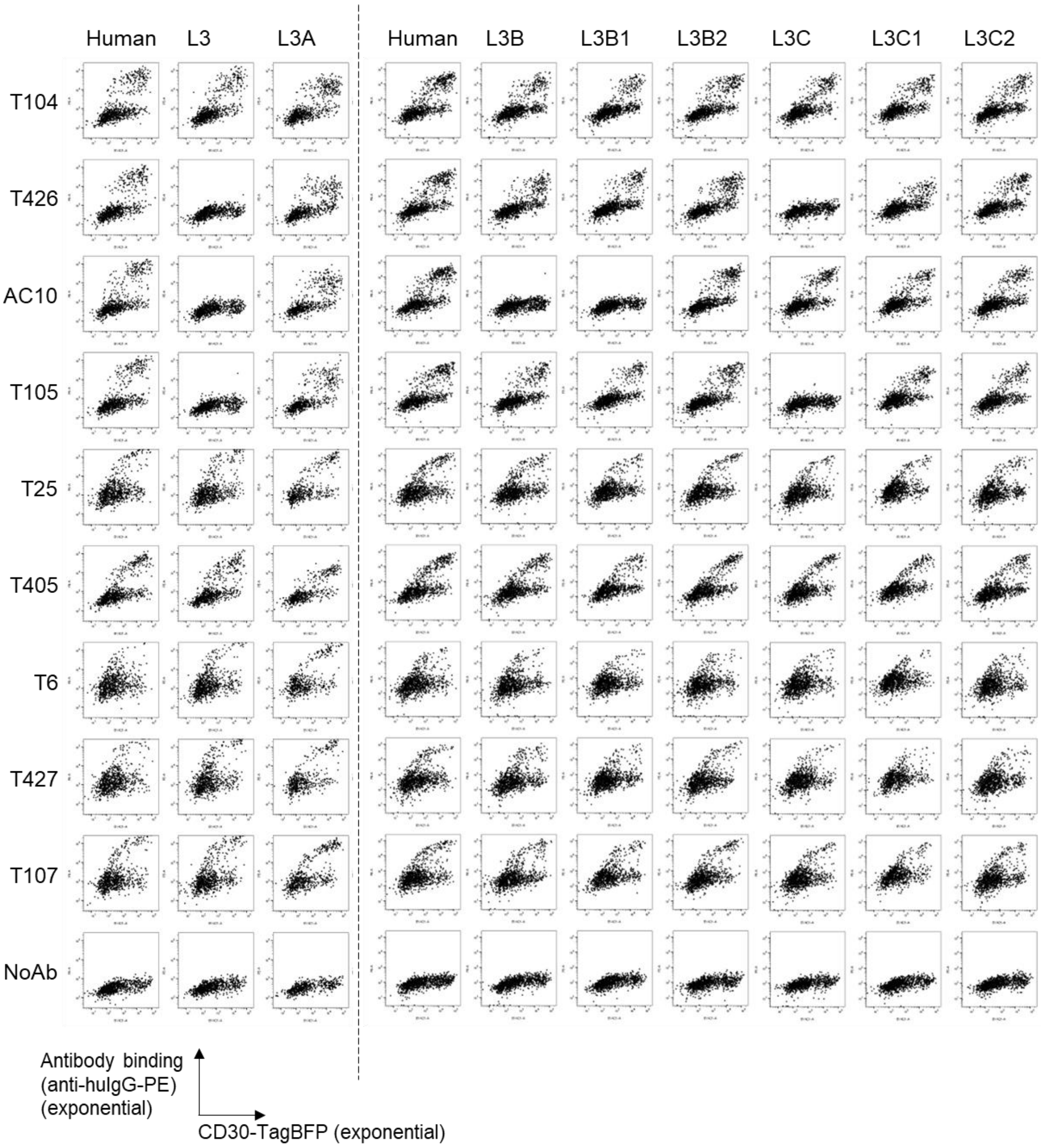


**Fig. S6.** Antibody binding to human CD30 with a portion of CRD3 substituted with L.pard sequences. Split line indicates data from independent experiments.

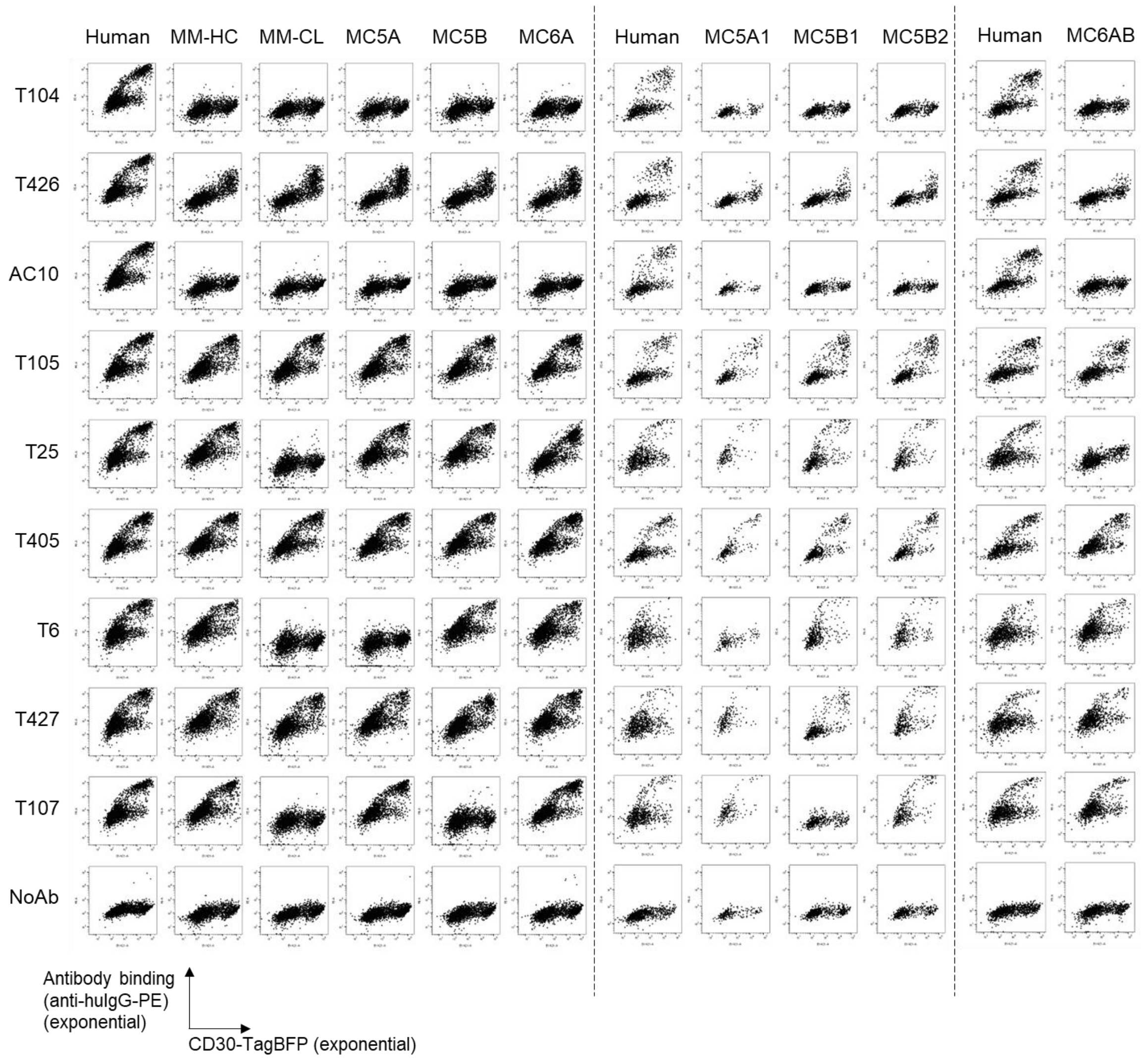


**Fig. S7.** Antibody binding to human CD30 with entire CRD1-3 substituted with M.myot sequence and a portion of CRD1 substituted with C.lani sequences. Split line indicates data from independent experiments.

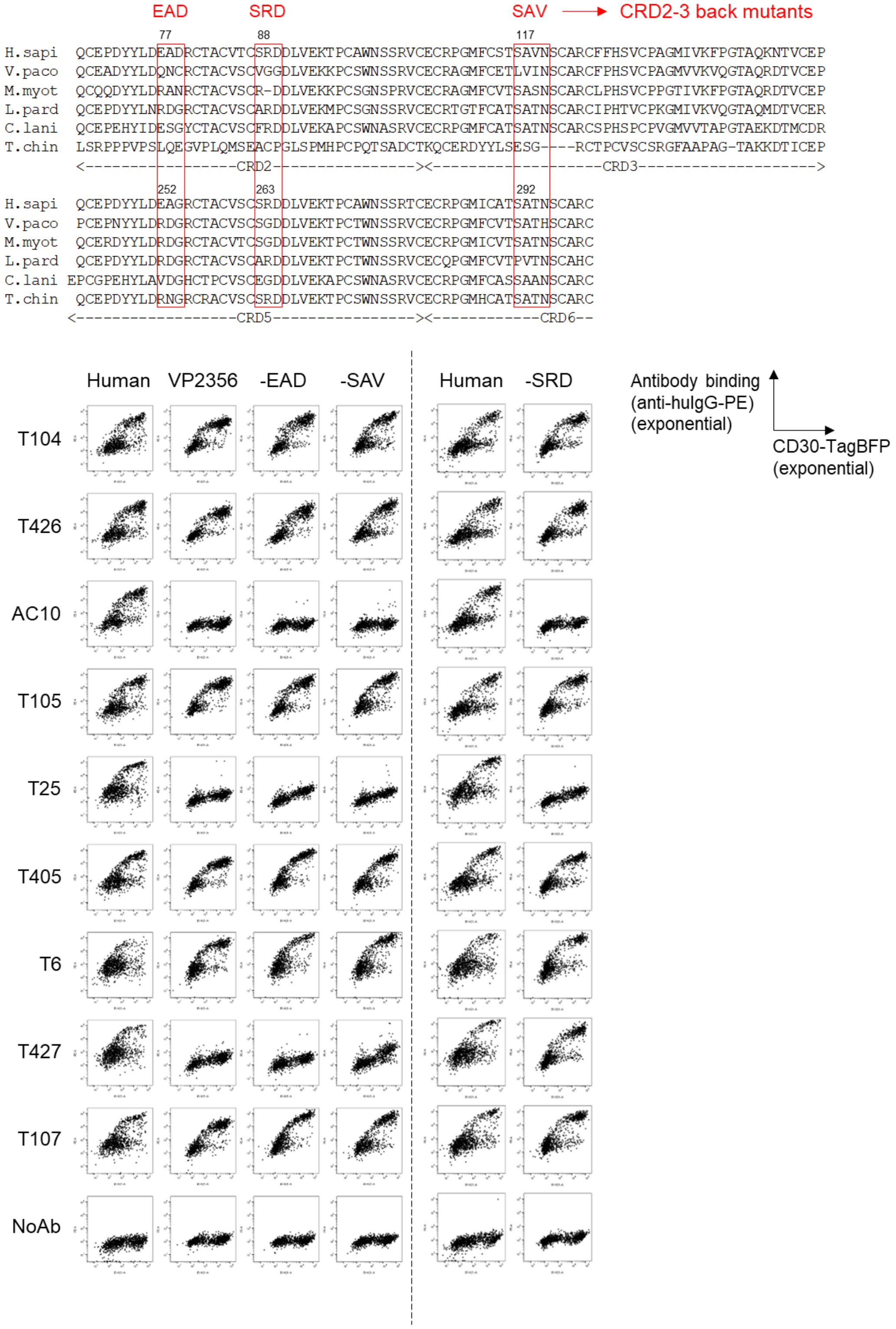


**Fig. S8.** Design of V.paco mutants and antibody binding to human CD30 with CRD2-3 and CRD5-6 substituted with V.paco sequences. "-EAD," "-SAV," and "-SRD" indicate back mutations to human sequences. Split line indicates data from independent experiments.

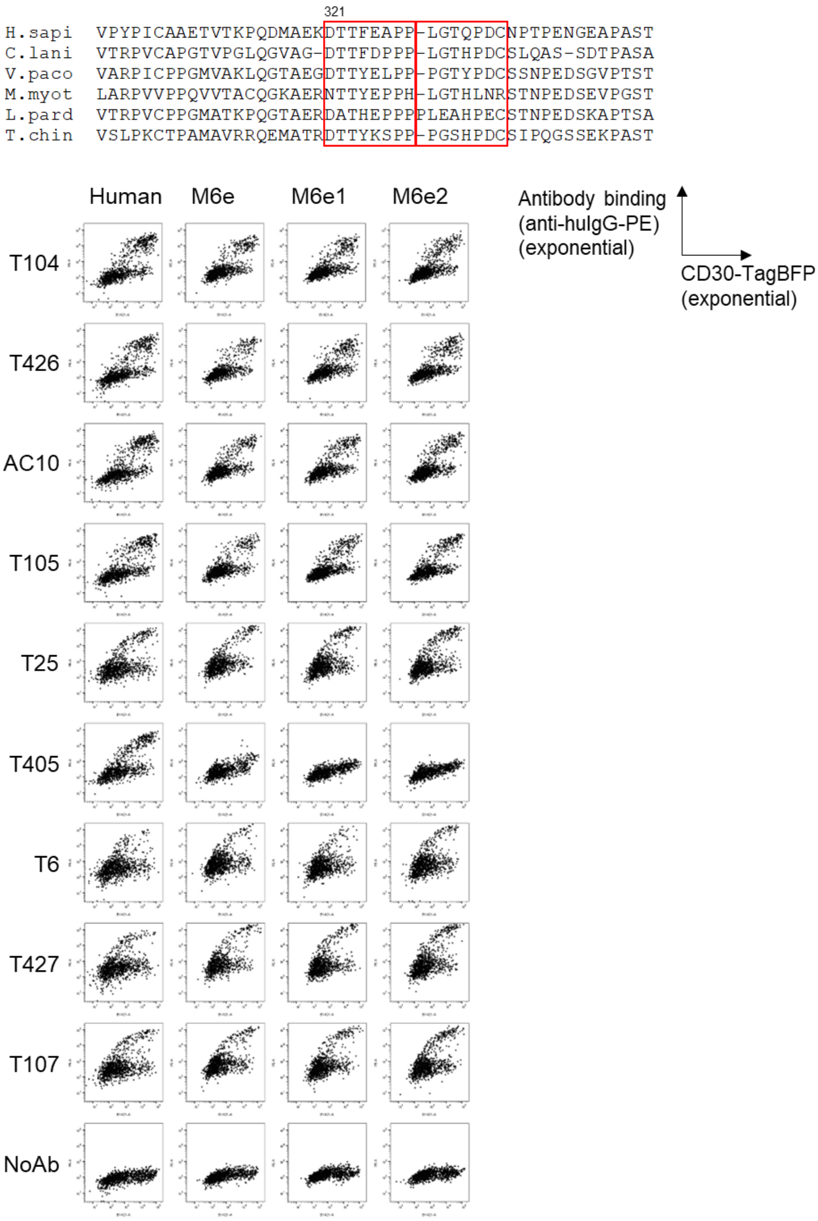


**Fig. S9.** Design of M.myot mutants and antibody binding to human CD30 with a portion of CRD6 or its C-terminal region substituted with M.myot sequences.

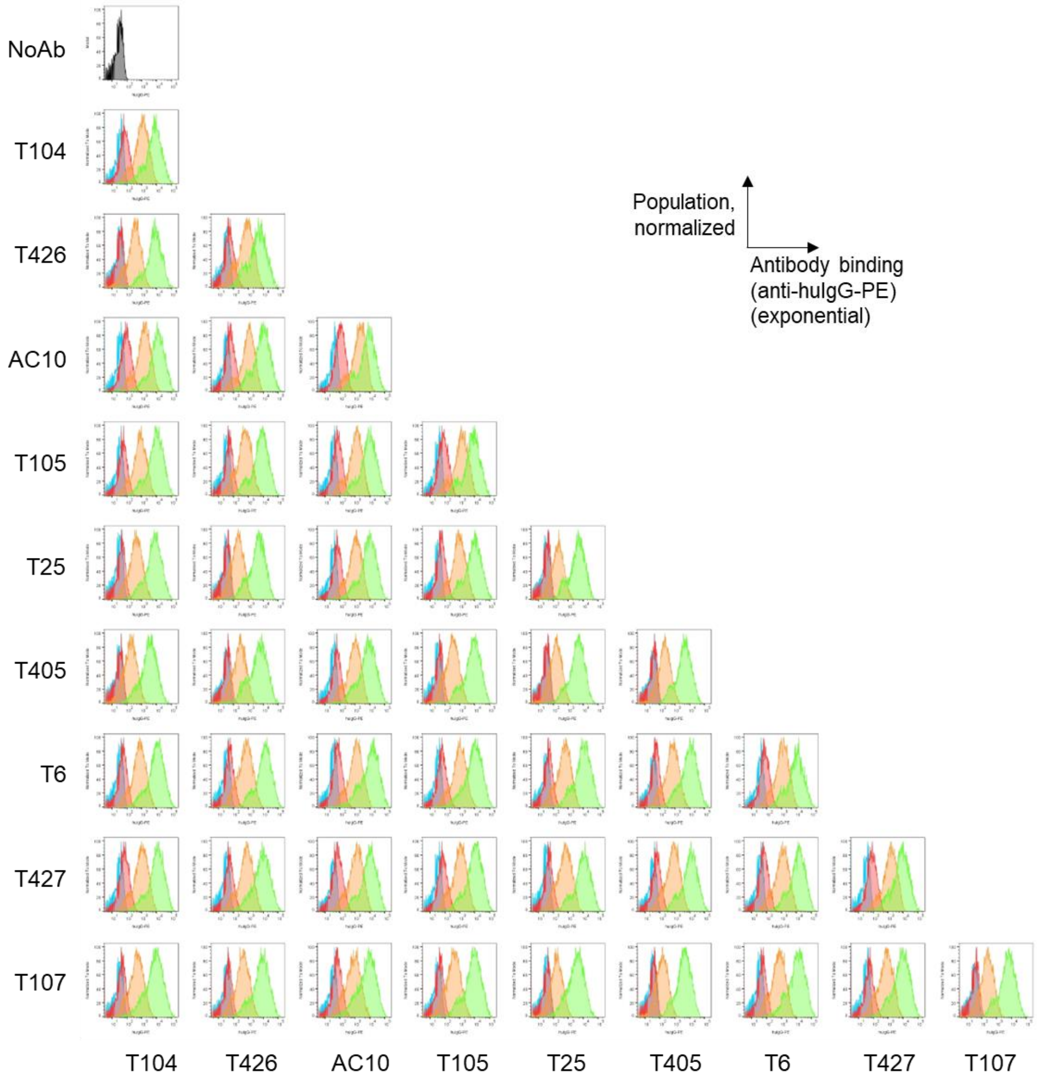


**Fig. S10.** Antibody binding to CD30-transfected Ramos-blue cells at various concentrations. Antibodies were tested at concentrations of 1500 ng/mL (green), 15 ng/mL (orange), 0.15 ng/mL (red), or 0.0015 ng/mL (cyan). Each panel corresponds to a specific antibody configuration: when x- and y-axis labels are the same (e.g., both are T104), it represents the binding of cAb (e.g., T104). Conversely, when x- and y-axis labels are different (e.g., x-axis label is T104 and y-axis label is T426), it signifies the binding of BpAb with the labeled Fvs (e.g., BpT104-426).

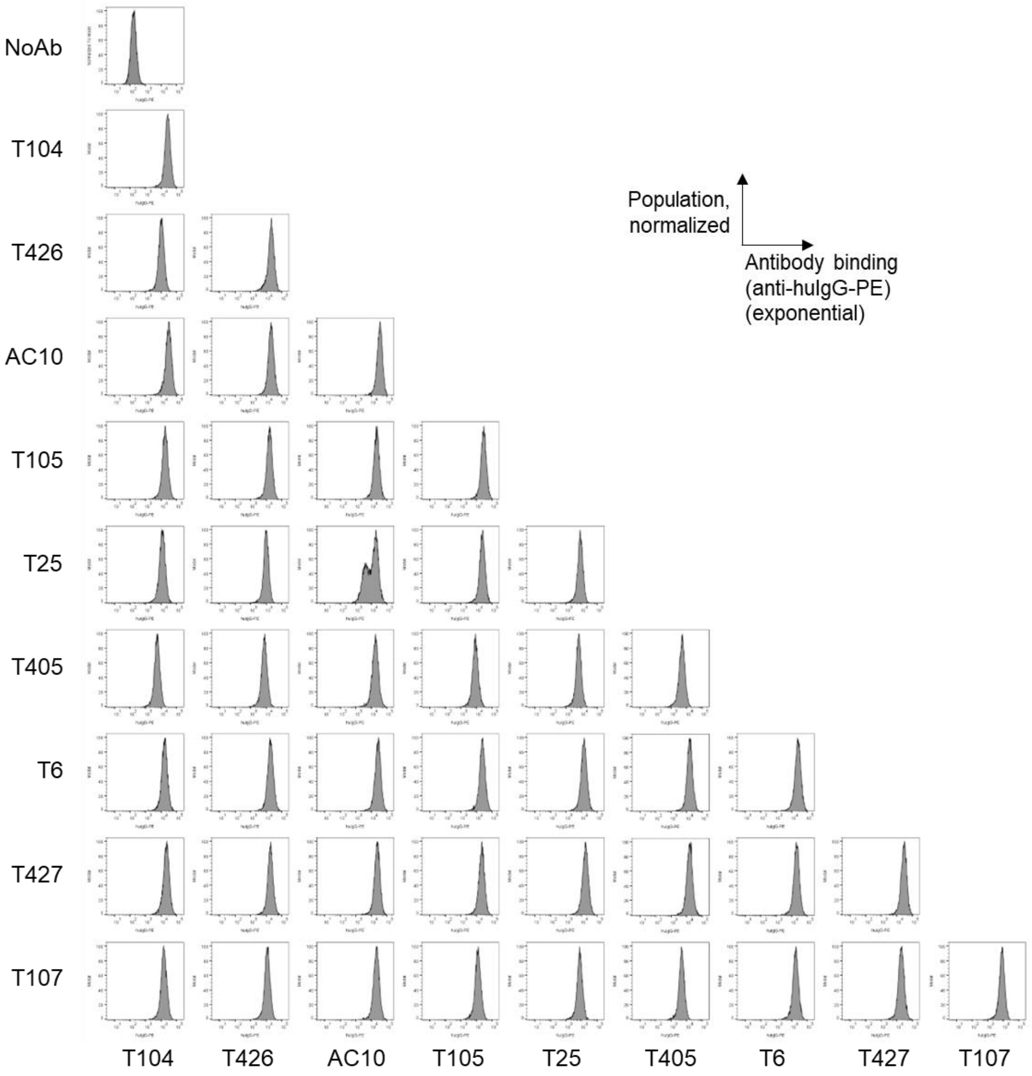


**Fig. S11.** Antibody binding to KARPAS 299 cells at 150 ng/mL. For each panel, when x- and y-axis labels are the same (e.g., both are T104), the panel corresponds to binding of cAb (e.g., T104). When x- and y-axis labels are different (e.g., x-axis label is T104 and y-axis label is T426), the panel corresponds to binding of BpAb with labeled Fvs (e.g., BpT104-426).

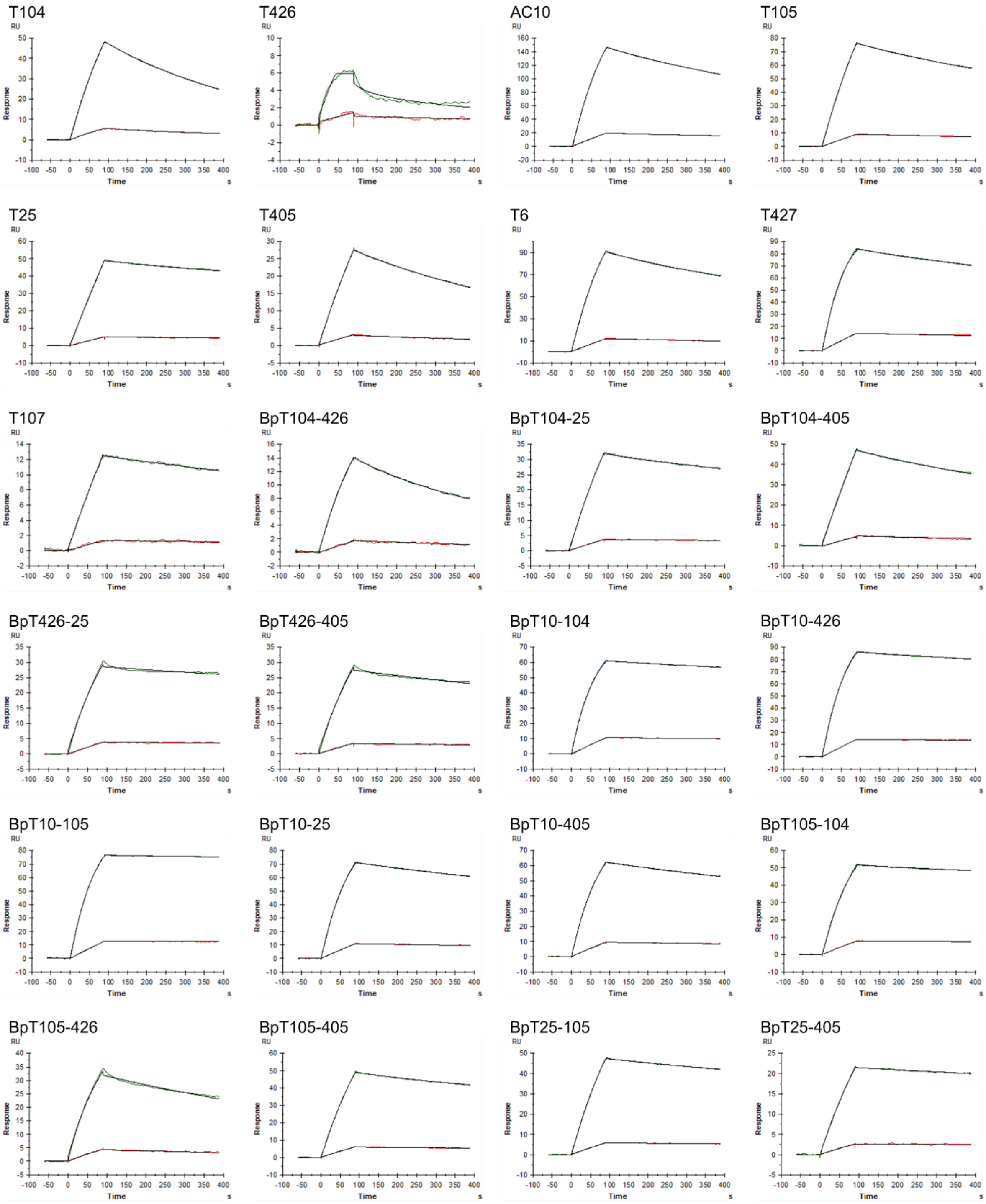
T104
T426
AC10
T105
T25
T405
T6
T427
T107
BpT104-426
BpT104-25
BpT104-405
BpT426-25
BpT426-405
BpT10-104
BpT10-426
BpT10-105
BpT10-25
BpT10-405
BpT105-104
BpT105-426
BpT105-405
BpT25-105
BpT25-405
RU
Response
Time
s

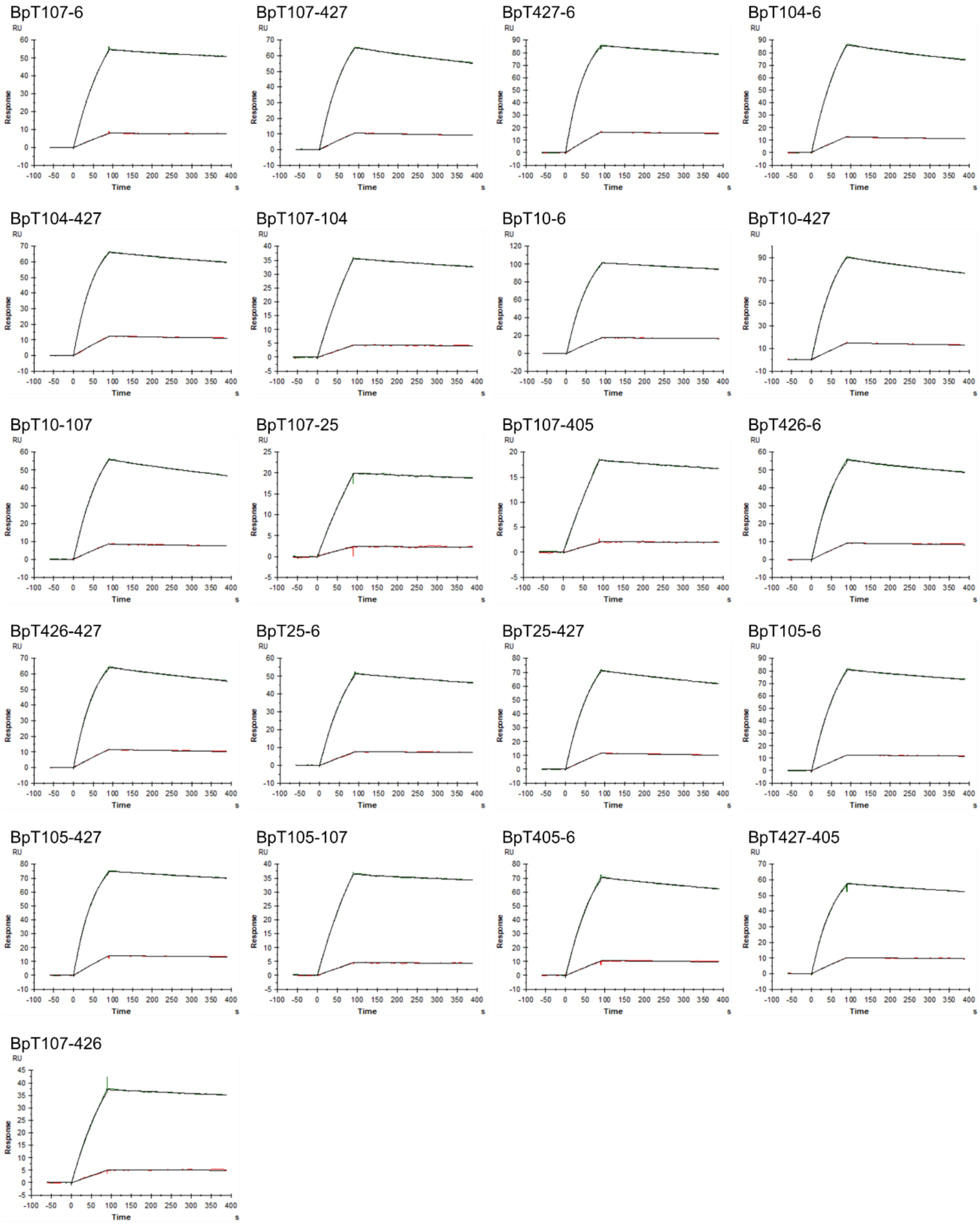


**Fig. S12.** Surface plasmon resonance charts of CD30-MBP binding to immobilized antibodies. CD30-MBP was flowed at concentrations of 2 nM (red) or 20 nM (green) for all antibodies except T426. The sensorgrams are displayed with the baseline subtracted for CD30-MBP flowed at 0 nM. For T426, the concentrations were 20 nM (red) or 200 nM (green). Black lines represent the fitting curves for a 1:1 binding mode.

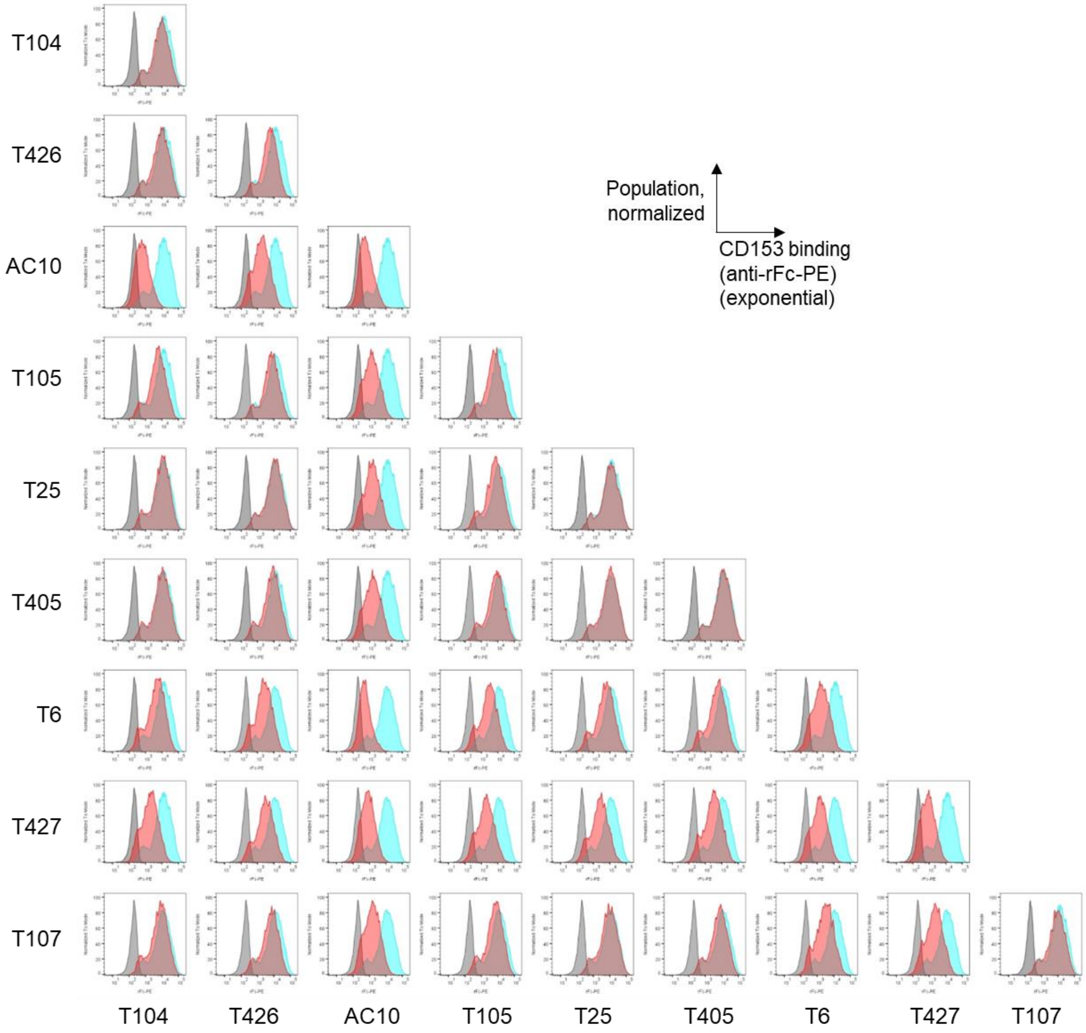


**Fig. S13.** Binding inhibition of CD153-rFc by cAbs and BpAbs. The binding of CD153-rFc to CD30-RamosBlue cells was assessed using flow cytometry. In the graphs, grey represents the secondary antibody (anti-rabbit Fc-PE) only, cyan indicates CD153-rFc alone, and red shows CD153-rFc in the presence of the antibody. Each panel corresponds to either cAb competition (e.g., T104) or BpAb competition with labeled Fvs (e.g., BpT104-426), depending on the alignment of the x- and y-axis labels.

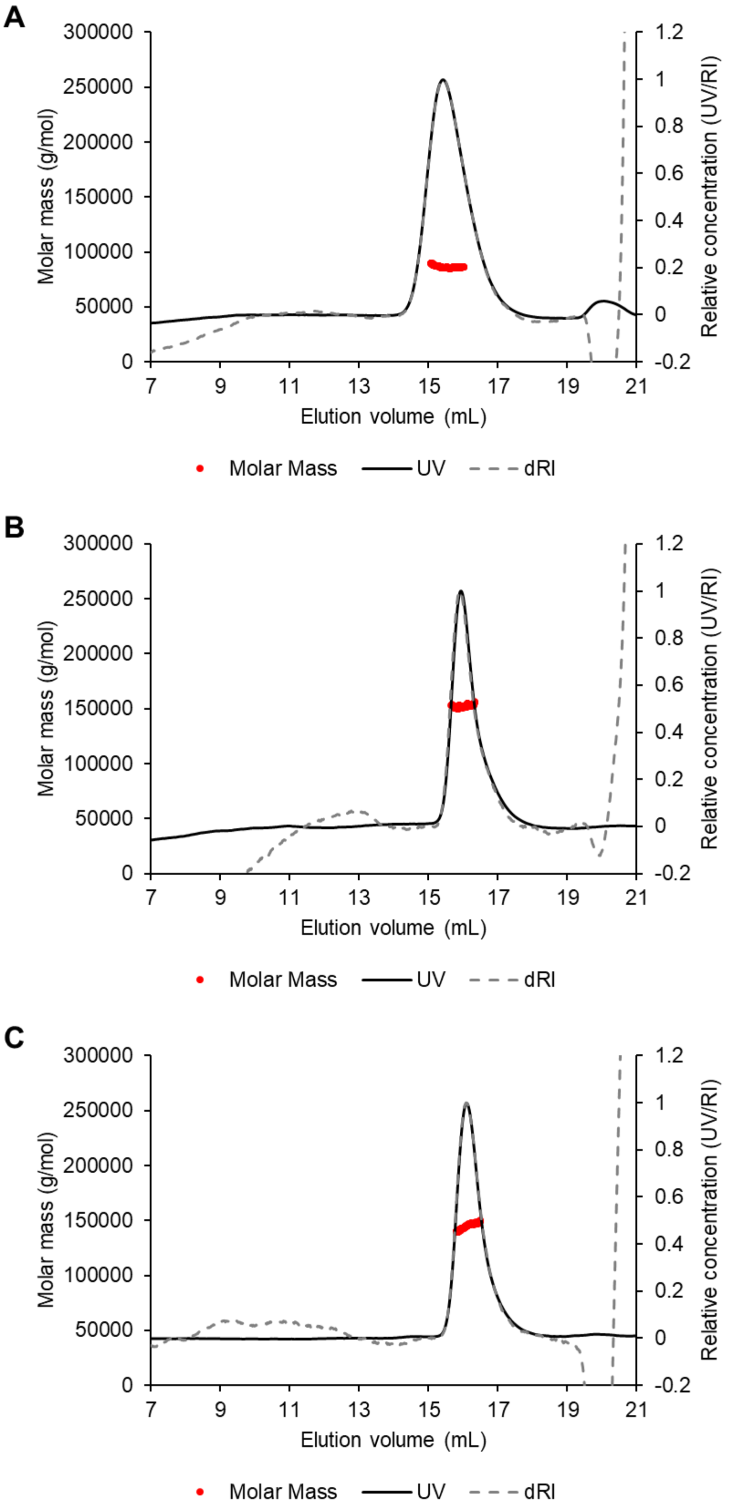


**Fig. S14.** SEC-MALS charts for individual components. Panels display the results for CD30-MBP alone (A), BpT10-104 alone (B), and BpT10-405 alone (C), each analyzed separately outside of the complex.
